# The necessity of considering enzymes as compartments in constraint-based genome-scale metabolic models

**DOI:** 10.1101/2022.12.14.520512

**Authors:** Xue Yang, Zhitao Mao, Jianfeng Huang, Ruoyu Wang, Huaming Dong, Yanfei Zhang, Hongwu Ma

## Abstract

As the most widespread and practical digital representations of living cells, metabolic network models have become increasingly precise and accurate. By integrating cellular resources and abiotic constraints, the prediction functions were significantly expanded in recent years. However, we found that if unreasonable modeling methods were adopted due to the lack of consideration of biological knowledge, the conflicts between stoichiometric and other constraints, such as thermodynamic feasibility and enzyme resource availability, would lead to distorted predictions. In this work, we investigated a prediction anomaly of EcoETM, a constraints-based metabolic network model, and introduced the idea of enzyme compartmentalization into the analysis process. Through rational combination of reactions, we avoid the false prediction of pathway feasibility caused by the unrealistic assumption of free intermediate metabolites. This allowed us to correct the pathway structures of L-serine and L-tryptophan. Specific analysis explains the application method of EcoETM-like model, demonstrating its potential and value in correcting the prediction results in pathway structure by resolving the conflict between different constraints and incorporating the evolved roles of enzymes as reaction compartments. Notably, this work also reveals the trade-off between product yield and thermodynamic feasibility. Finally, we provide a preliminary comparison of the thermodynamic feasibility of ammonia and glutamine as amino donors, which revealed that the direct utilization of ammonia does not have a decisive impact on the thermodynamic feasibility of the anthranilate pathway. Our work is of great value for the structural improvement of constraints-based models.

## 1. Introduction

With the enhancement of parameter acquisition ability and the improvement of functional annotation, the available data has experienced a blowout growth (Du et al., 2018b; Flamholz et al., 2011; Heckmann et al., 2018; Joshi et al., 2021; Li et al., 2022). For the construction of genome-scale metabolic models (GEMs), the most instructive digital method of cell metabolic processes was developed more than 20 years ago(Gu et al., 2019; Seif and Palsson, 2021), and the desire for refinement is becoming more ambitious than ever before. The automated construction process(Lu et al., 2019; Machado et al., 2018; Seaver et al., 2021) and quality control methods(Lieven et al., 2020) of metabolic network models have gradually become mature(Mendoza et al., 2019). In recent years, enzyme-constrained models were developed continuously, and automated processes developed based on the MOMENT principle(Adadi et al., 2012), such as GECKO(Sanchez et al., 2017), sMOMENT(Bekiaris and Klamt, 2020) and ECMpy(Mao et al., 2022), have been migrated rapidly(Ye et al., 2020). At the same time, enzyme-constrained models have been successfully integrated with thermodynamic constraints, such as ETFL (metabolism and expression model framework with thermodynamic constraints) (Salvy and Hatzimanikatis, 2020) and ETGEMs (GEMs integrating enzymatic and thermodynamic constraints)(Yang et al., 2021).

During ETGEM development, we realized that due to the excessive separation of some chemical reactions in the *i*ML1515 stoichiometric framework(Monk et al., 2017), some prediction results were inconsistent with reported facts after the integration of thermodynamic constraints. For example, the combination of PGK_r, PGCD, GAPD, TPI and FBA reactions (catalyzed by phosphoglycerate dehydrogenase, phosphoglycerate kinase, glyceraldehyde 3-phosphate dehydrogenase, fructose-bisphosphate aldolase and triose-phosphate isomerase, respectively) was mistakenly considered to be thermodynamic infeasible(Yang *et al*., 2021). Therefore, it is time to split and merge GEM reactions according to the real intracellular situation. We also realized that if the biochemical reactions were excessively split, the correct understanding of remarkable adaptive strategies that cells have evolved, such as enzyme complexes, fusion enzymes and multifunctional enzymes, would be missed(Hwang and Lee, 2019; Martínez et al., 2014; Skirgaila et al., 2013). Conversely, if we take the initiative to disassemble the links between reactions, we may obtain new possibilities in pathway diversity, broaden the solution space, and develop new compounds synthesis method.

The concept of compartmentalization is frequently mentioned both in model construction and practical research. Prokaryotic GEMs are generally divided into two or three regions including cytoplasmic, extracellular and periplasmic compartments, whereby the latter is only present in Gram-negative bacteria, while conditional connectivity is achieved through transport and exchange reactions. In GEMs of eukaryotes such as of *Saccharomyces cerevisiae* and *Yarrowia lipolytica*, here are more than ten independent cell divisions such as mitochondria and other organelles(Förster et al., 2003). The compartmentalization of engineered metabolic pathways in yeast mitochondria notably improved the production of branched-chain alcohols (Avalos et al., 2013; Zhang et al., 2022; Zhang et al., 2019). Dating back further, the concept that enzymes and their microenvironment should be considered as compartments was proposed(Srere and Mosbach, 1974). Unfortunately, such a microscopic compartmentalization concept has not been effectively and clearly absorbed in the construction and application of metabolic network models or their extended versions.

In this work, we used the model EcoETM(Yang *et al*., 2021), an extended *i*ML1515 by integration of thermodynamic and enzymatic constraints. By analyzing the synthetic pathways of L-serine and L-tryptophan, we revealed the necessity of considering the evolutionary strategy of multifunctional enzymes and enzyme complexes as metabolic compartments to effectively avoid unreasonable conflicts between different constraint levels in the extended GEMs. Finally, we demonstrated the application method and value of the multiple-constraint GEMs in the analysis of product synthesis pathways and improving the basic framework quality of GEMs.

## 2. Materials and Methods

The concentration boundaries for metabolites, including O_2_ (0.5 - 200 μM)(Murphy, 2009; Reynafarje et al., 1985) and CO_2_ (0.1 - 100 μM)(Hadicke et al., 2018), were are the same published before(Yang *et al*., 2021). In addition, because high concentrations of ammonia are toxic(Javelle et al., 2004; Muller et al., 2006), the ammonia concentration is set to a more realistic range of 10 μM - 1 mM(Kim et al., 2012). All methods of the reaction processing and parameter settings were listed in the corresponding positions of the manuscript.

All of the simulations were performed in Jupyter notebook with ETGEMs, a Python-based tool for multi-constrained metabolic network model construction adopting the Cobrapy toolbox(Gollub et al., 2021) and Pyomo software package. Codes and the EcoETM model are available at https://github.com/tibbdc/ETGEMs.

## 3. Results

### 3.1 Prediction of the serine synthesis pathway using EcoETM

In our previous publication(Yang *et al*., 2021), we identified five distributed bottleneck reactions(Mavrovouniotis, 1993), PGCD, PGK_reverse, GAPD, FBA and TPI (see table S1 of this manuscript and in attached table F of the cited reference), which led to the false prediction that the pathway in which they participate jointly is thermodynamically infeasible. Among them, PGCD is a well-known reaction in the L-serine synthesis pathway, and the other four reactions are also central to the glycolysis pathway. This subset is suitable as an example to illustrate the concept of distributed bottleneck reactions, but the conclusion that the classical EMP pathway cannot coexist with the conventional L-serine synthesis process is inconsistent with observations. Therefore, we analyzed the serine synthesis pathway in detail. The maximal thermodynamic driving force (MDF) of the L-serine synthesis pathways is variable, with 13 turning points, as shown in Fig. 1. The information of the bottleneck reaction(s) causing the MDF decline at each turning point is shown in table S1, and the corresponding pathways at the turning points are detailed in Fig. S1.

**Fig. 1.**
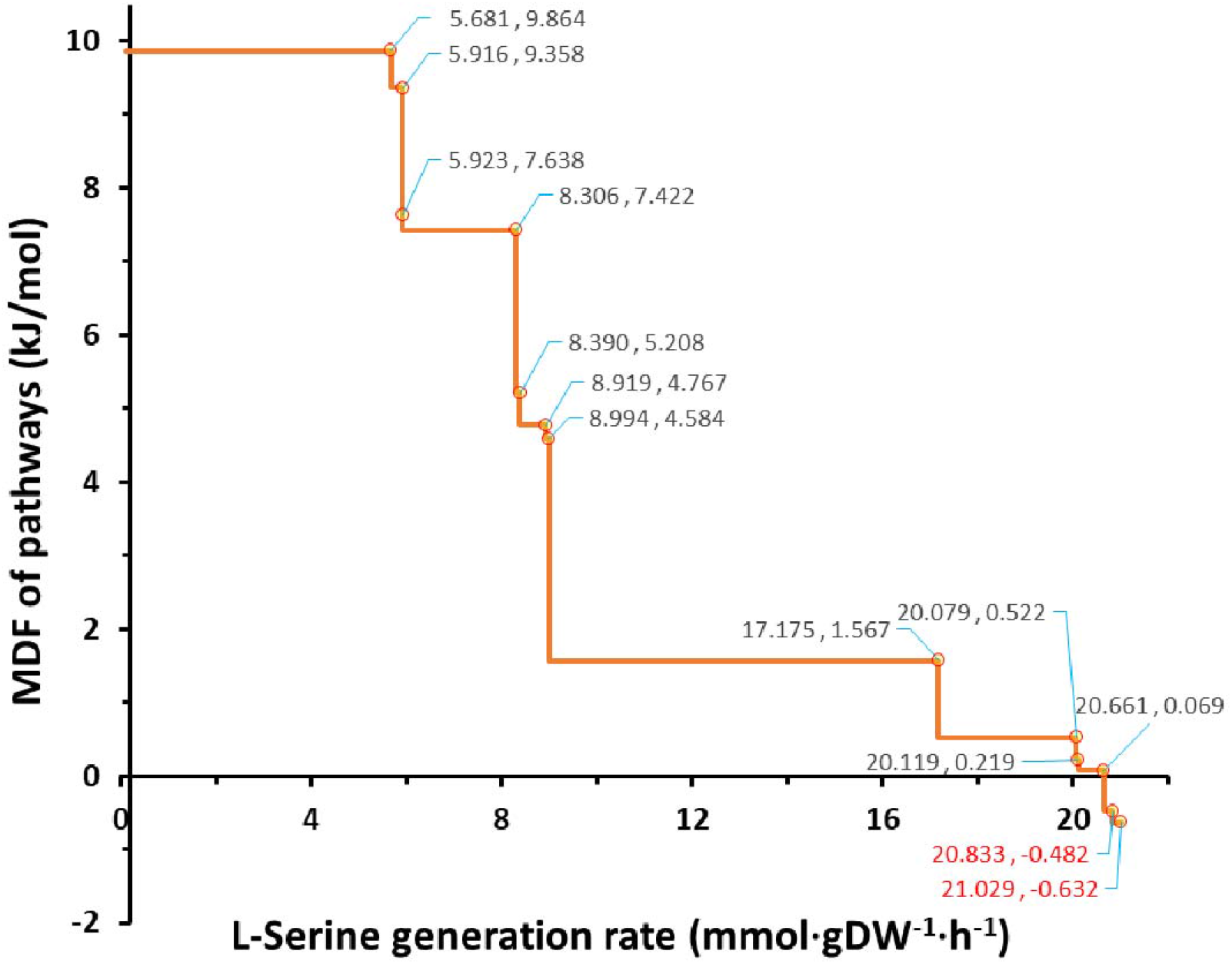
The maximal thermodynamic driving force (MDF) of the serine synthesis pathway. The horizontal coordinate value indicates the rate of L-serine synthesis, the grey value at the turning point indicates the MDF level of pathway(s). The upper bound of the glucose uptake rate was set to 10 mmol/g DW/h.

The process of serine synthesis is divided into 13 stages according to the MDF levels. The MDF level in the first seven stages can reach more than 4 kJ/mol, and the thermodynamic feasibility is ideal. The MDF level of the middle four stages is close to zero, and the feasibility is quite low. Finally, the MDF of the last two stages is negative, so it can be considered that they are thermodynamically infeasible.

Further analysis revealed that there are 2 localized and 11 distributed bottleneck reactions in total, representing the combination of reactions located in the central metabolic pathways (mainly in EMP, PPP and TCA)(Orsi et al., 2022). For the yield indicator, the highest flux level that can be achieved at the end of the seventh stage is 8.99 mmol/gDW/h, which is only 42.7% compared with the highest flux of 21.03 mmol/gDW/h, and is unsatisfactory since it represents 57% carbon loss.

From the eighth stage, the MDF fell to 1.57 kJ/mol, which was caused by the localized bottleneck reaction of PGCD alone. Although the participation of this reaction reduced the MDF level of the pathway(s), it contributed significantly to the increase of yield, so that the synthetic flux of the pathway(s) reached 17.175 mmol/gDW/h, which was equivalent to 81.7% of the maximal yield.

From the ninth to the twelfth stage, the thermodynamic feasibility of the pathway(s) decreased gradually, which was caused by several distributed bottleneck reactions. The common point of these combinations of distributed bottleneck reactions is that the PGCD reaction is always sharing limiting metabolites with other bottleneck reactions, the overall thermodynamic feasibility of the pathway(s) is lower than in the eighth stage (when PGCD is used as a localized bottleneck reaction). According to the second law of thermodynamics (MDF≥0), the flux limit of the last thermodynamically feasible pathway(s) at the end of the eleventh stage is 20.66 mmol/gDW/h, which is equivalent to 98.2% of the maximal yield.

At the thirteenth stage, there are still three distributed bottleneck reactions, but PGCD is not among them. By investigating the individual thermodynamic driving force levels of the three reactions, it was found that they are thermodynamically feasible in principle (in table S2). However, due to the close association in sharing limiting metabolites, their simultaneous activity can make the pathway thermodynamically infeasible. This also shows that the thermodynamic feasibility based on the flux balance analysis (FBA) in GEMs with integrated multi-constraints is different from the simple preset reaction direction and reversibility. It should be noted that some anaerobic organisms, such as *Desulfovibrio desulfuricans*(Sánchez-Andrea et al., 2020) and *Clostridium drakei*(Song et al., 2020), adopt the POR5_r reaction (catalyzed by pyruvate synthase) to implement the reductive glycine pathway for autotrophic CO_2_ fixation(Bar-Even et al., 2011). In addition to POR5_r, FLDR2 (catalyzed by flavodoxin reductase) is the only reaction that can reduce the semi-oxidized flavodoxin in the *i*ML1515 model. As a reaction that consumes reduced flavodoxin, POR5_r inevitably shares fluxes with FLDR2. In addition, FLDR2 also shares the b0684 and b2895 genes (*fldA* and *fldB*), encoding flavodoxin 1 and 2 respectively, with the POR5_r reaction. The coupling relationship between FLDR2 and POR5_r has been described by Maurice et al (St Maurice et al., 2007). Recently, a similar coupling process has been used to construct an artificial minimal carbon fixation pathway, but the difference is only that several different ferredoxins were used as the electron donor instead of flavodoxin(Xiao et al., 2022). Similar to the above two reactions, the PFL reaction is also only effective under anaerobic conditions (Wang et al., 2010), and requires the activation of flavodoxin reductase (FLDR2)(Ingelman et al., 1997). The three reactions can form a pathway under anaerobic conditions and the net reaction is the synthesis of formate from CO_2_, consuming a reducing equivalent provided by NADPH (table S4). For *E*.*coli*, formate is the necessary source of reducing power in anaerobic deoxyribonucleotide synthesis(Mulliez et al., 1995).

Based on these results, it can be found that the PGCD reaction has a strong influence on the L-serine yield. With its participation, the maximum yield of pathway(s) can be increased from 42.7% to 98.2% of the theoretical yield predicted by the initial GEM, *i*ML1515. According to the MetaCyc database, the PGCD reaction should indeed appear in the L-serine synthesis pathway of *E. coli*, which was proved by experimental results (in Figure S2).

In the combination of bottleneck reactions in stages 9-13 (table S1), the distributed bottleneck reactions with or without PGCD are always located outside the core L-serine synthesis pathway (see Figure S2 and Table S2). This indicates that the thermodynamic analysis and optimization of specific pathways alone may be inapplicable when they are placed in the cellular context, which reflects the differences between the solution for pathway MDF in the network(Hadicke *et al*., 2018) and the thermodynamic evaluation for specific pathways(Noor et al., 2014), indicating that the thermodynamic evaluation of preset pathways is likely to misdiagnose the actual situation in the complex intracellular system.

### 3.2 Learning the strategy from cells to overcome the thermodynamic bottleneck

The PGCD reaction is important for producing a high yield of L-serine, but its unfavorable thermodynamics are also a problem that must be solved. As shown in Fig. 2A, the PGCD reaction will release the reducing force of NADH. In 2017, Zhang et al. (Zhang et al., 2017) proved that when PGCD is coupled with the reduction force consumption reaction catalyzed by the same enzyme expressed by the identical gene, there is a significant improvement of its thermodynamic feasibility.

**Fig. 2.**
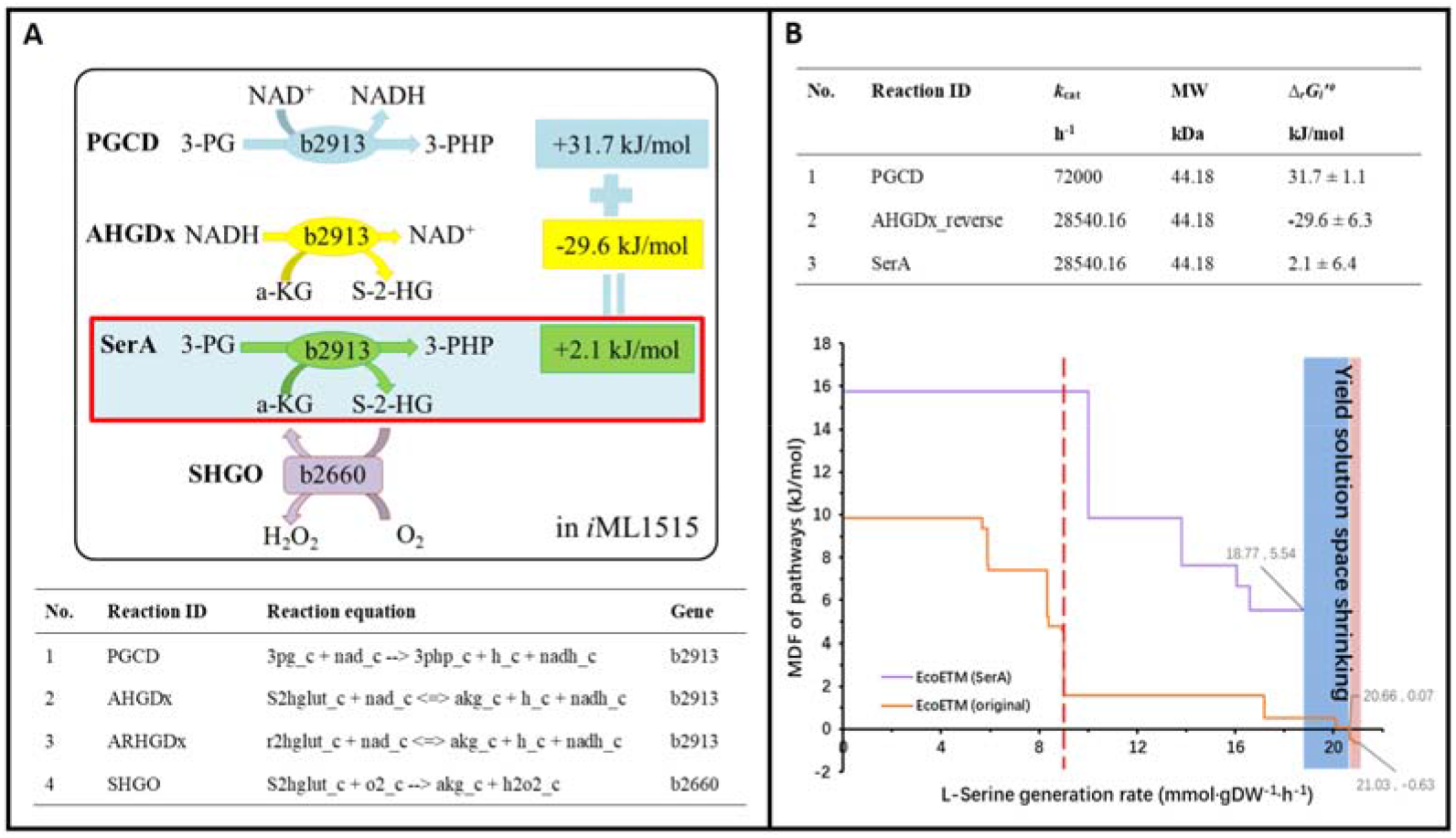
Strategy for resolving the thermodynamic bottleneck in L-serine synthesis pathways. A. The biological phenomena and principles reproduced in this figure refer to the work performed by Zhang et al. (Zhang *et al*., 2017). B. Thermodynamic feasibility analysis and comparison of L-serine synthesis processes. The result predicted by the initial model is indicated by the orange line, and the result predicted by models with combined reaction is indicated by the purple line. The left side of the red dotted line is the high thermodynamic feasibility range predicted by the initial model, while the shaded part represents the reduction of the yield space, while the blue part represents the space reduction of the thermodynamically feasible yield.

In the *i*ML1515 model, there are two potential natural coupling reactions with PGCD, AHGDx and ARHGDx (Fig. 2A). The above three reactions share the identical gene number (No. b2913), which therefore encodes a multifunctional enzyme. In the initial *i*ML1515 model, the AHGDx and ARHGDx reactions were reversible, with the only difference between them being the substrate configuration. However, in the whole metabolic network, there is neither the configuration conversion process between their substrates, nor the consumption or silencing reaction with *R*-configuration substrate (*R*-2-hydroxyglutate, r2hglut), so ARHGDx is redundant and cannot participate in flux balance analysis. The *S*-configuration substrate of the AHGDx reaction (*S-*2-hydroxyglutarate, s2hglut) can only be used as the substrate of another irreversible reaction, SHGO, namely there are no other means to consume it. Therefore, it can be considered that the positive reaction direction of AHGDx in the model is also redundant, and only its reverse reaction is effective. The AHGDx_reverse reaction consumes reducing force, so when it is coupled with the SHGO reaction, the generation and consumption of carbonaceous metabolites can also be offset. This means that when the combined reaction continues to be coupled with PGCD, the thermodynamic feasibility of the overall reaction can be promoted by balancing the reducing force, and the metabolite 2-oxoglutarate (*α*-ketoglutarate, *α*-KG) can also be supplemented, as shown in Fig. 2A.

It is worth noting that the coupling of reactions is often used in enzyme activity detection(Cui et al., 2018), the construction of multi-enzyme system(Cai et al., 2021; Schoffelen and van Hest, 2013) and thermodynamic optimization of pathways(Lin et al., 2018; Yang et al., 2019), but the synthesis process of L-serine revealed by Zhang et al. (Zhang *et al*., 2017) is fundamentally different. The main reason is that it is not only based on the coupling between reactions to change the metabolites concentrations in the cell, but there is also a correlation at the genetic level, so as to realize the reaction coupling within a protein structure, which offers a natural advantage for the transmission and capture of metabolites between reactions to strongly increase the thermodynamic driving force. This illustrates the significance of enzyme complexes and multifunctional enzymes, which can be considered the most basic level of cellular compartmentalization(Kojima and Takayama, 2018).

Then, a combined reaction catalyzed by SerA with a Δ_*r*_*G*_*i*_′[ of 2.1 □/mol was added to the model, and the equation was set as “3pg_c + akg_c --> 3php_c + S2hglut_c”, which was adopted to replace the original two partial reactions PGCD and AHGDx_ reverse. Since the catalytic efficiency of enzymes is limited by the reaction with the poorest kinetic parameters, after comparing the *k*_cat_ of PGCD and AHGDx_reverse, the inferior *k*_cat_ of the AHGDx_reverse reaction was assigned to the combined reaction SerA, as shown in Fig. 2B.

After the combination of reactions, the change of MDF in the L-serine synthesis process was recalculated (as shown in Fig. 2B). The pathway thermodynamic driving force was significantly improved. At the stage where the original thermodynamic feasibility reaches a good level (left side of the red dotted line), the achievable flux increased from 8.99 to 18.77 mmol/gDW/h. Moreover, after correction by merging reactions, the thermodynamic driving force in the whole yield space was optimized. Due to the double adjustment of reaction stoichiometric and enzyme kinetic parameters, the yield solution space was reduced by 9.1% (relative to the original thermodynamic feasible yield solution space, blue shaded area) to 10.7% (relative to the original total yield solution space, whole shaded area) as shown in Fig. 2B. In addition, the number of MDF levels was reduced and the pathway was only divided into five stages. See Fig. 2B and attached table S3.

Because the POR5_r, FLDR2 and PFL reactions only work under anaerobic conditions, we closed the oxygen exchange reaction and combined the three reactions into one reaction named POFLx (table S4-6), and then predicted the L-serine synthesis pathway(s). As shown in table S7 and Fig. S3, only three MDF levels were predicted, and the yield of the three pathways was generally low, generating by-products such as acetate, ethanol, lactate and DHA. In previous studies, overflow metabolism under aerobic conditions was attributed to the limitation of enzyme resources, which did not reach the upper limit (<0.13 g enzyme/gDW) in the simulation results under anaerobic conditions. Therefore, the formation of fermentation products under anaerobic conditions is mainly caused by topological (for stages 1 and 2) and thermodynamic (for stage 3) reasons. Unsurprisingly, the absence of oxygen and the reactions it participates in greatly reduces the pathway solution space.

Among these three pathways, only the pathway of the third stage with the highest yield adopted the combined POFLx reaction, and the thermodynamic level of the pathway has not been improved at all. The MDF of the combined POFLx reaction was −1.897 kJ/mol, the sum of the MDF values (−0.632 kJ/mol) of the original three partial reactions. Then, we listed the optimal concentration levels of all metabolites involved in the three partial reactions (table S8) and the combined reaction (table S9). It can be seen that the intermediate metabolites avoided by the reactions combination are all within the concentration boundaries, while the NADPH / NADP (substrate, concentration ratio is limited to a narrow range), limiting metabolites CO_2_ (substrate, low solubility) and formate (product) are all retained in the combined reaction. On the one hand, the thermodynamic feasibility of the partial reaction is overestimated due to the lack of a concentration ratio constraint of reduced and semi-oxidized flavodoxin in the model. Notably, it is especially easy to obtain this unfair advantage when the concentration constraint is relaxed in a multi-substrate reaction such as POR5_r. On the other hand, the enzyme conformation change of PFL involves the participation of high-energy S-adenosylmethionine (SAM)(Shisler et al., 2017), so that when only the energy levels of substrates and products are considered, the reaction thermodynamics is underestimated. When combined, these complex factors make it difficult to accurately evaluate the thermodynamic feasibility of the POFLx reaction.

### 3.3 Reanalysis of L-tryptophan synthesis pathways

L-Tryptophan is a synthetic precursor of many bioactive components with medicinal value(Chen and Zeng, 2017). As its essential precursor, the synthesis efficiency and feasibility of L-serine production should affect the performance of the L-Trp synthesis pathway. In previously published work, based on the unmodified EcoETM model of the L-serine synthesis process, the MDF levels of the L-tryptophan synthesis pathways were variable and always had acceptable thermodynamic feasibility (Fig. 3, the minimum value of MDF is 4.767 kJ/mol). This indicates that its pathway does not include PGCD reaction (the MDF is not greater than 1.57 kJ/mol when PGCD is involved). Hence, the recognized synthesis process of essential precursor L-serine was not adopted. Therefore, it was necessary to reanalyze the L-tryptophan synthesis process.

**Fig. 3.**
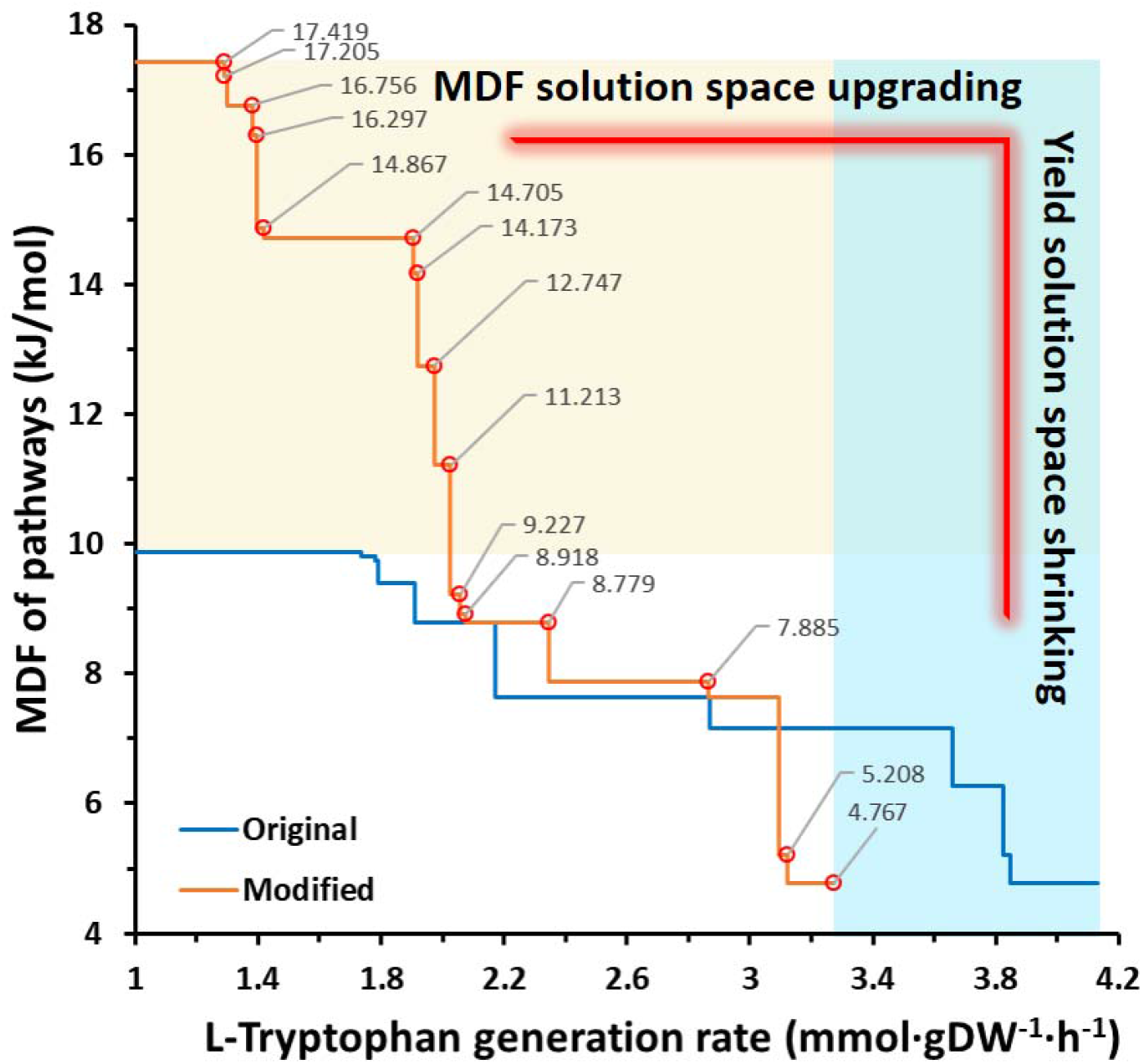
The MDF levels of the L-Trp synthesis pathways. Comparison of the MDF levels comparison of L-Trp pathways before (blue color) and after correction (orange color). The horizontal coordinate value indicates the L-Trp synthesis rate, and the grey value at the turning point indicates the MDF level of pathway(s). The upper bound of the glucose uptake rate was set to 10 mmol/gDW/h.

The initial MDF curve of the L-tryptophan synthesis pathway is shown in Fig. 3, and the analysis results of bottleneck reaction(s) at the turning point are shown in attached table S10.

In table S10, the MDF can be divided into ten levels. In the first level, the bottleneck reactions are the two incomplete semi-reactions ACONTa and ACONTb used to realize the production of isocitrate from citrate (via aconitate) in the TCA cycle. Based on the understanding of the partial reaction process catalyzed within the same whole enzyme structure, although aconitate is the product of the ACONTa reaction and the substrate of ACONTb, it is essentially different from the phenomenon of limited metabolite sharing between bottleneck reactions. Its generation and consumption are catalyzed by the same enzyme structure, which saves the time and energy costs for diffusion and capture, improving the affinity of the enzyme for the substrate molecule. By investigating the enzyme catalytic mechanism of aconitase, it can be found that the intermediate metabolite aconitate normally does not dissociate from the active site, so that almost none is released into the cytoplasmic environment(David L. Nelson and Cox, 2008; Morrison, 2021). Therefore, similar partial reactions should also be combined in the analysis process.

Most of the bottleneck reactions at stages 2 to 8 are found in central carbon metabolism, but there are also two additional high-frequency reactions, termed TRPS3 and TRPAS2_reverse. In the second stage, TRPS3 becomes one of the bottleneck reactions because it shares the metabolite G3P with the central metabolic reactions. As a direct synthesis reaction of L-tryptophan, TRPAS2_reverse often appears in pairs in the distributed bottleneck reactions from the third stage by sharing indole with the TRPS3 reaction. Contrary to expectations, L-serine is not required for TRPAS2_reverse to synthesize L-tryptophan, which directly leads to the absence of L-serine from the L-tryptophan synthesis pathway. Subsequently, we listed all the reactions in the *i*ML1515 model that may generate L-tryptophan directly, as shown in table 1.

**Table 1.** All reactions for direct L-Trp formation in the *i*ML1515 model

| No. | Reaction ID | Reaction equation | Gene ID | $\Delta_r G_i'$<br>kJ/mol |
| --- | --- | --- | --- | --- |
| 1 | TRPS1 | 3ig3p_c + ser__L_c --> g3p_c + h2o_c + trp__L_c | b1260 & b1261 | -30.5±6.2 |
| 2 | TRPS2 | indole_c + ser__L_c --> h2o_c + trp__L_c | b1260 & b1261 | -49.7±2.2 |
| 3 | TRPAS2_reverse | indole_c + nh4_c + pyr_c--> h2o_c + trp__L_c | b3708 | -21.1±1.1 |
| 4 | MTRPOX | Nmtrp_c + h2o_c + o2_c --> fald_c + h2o2_c + trp__L_c | b1059 | -44.6±13.7 |

All four reactions in table 1 had acceptable thermodynamic parameters, as illustrated by the Δ***rG***_***i***_′□values. Among them, both TRPS1 and TRPS2 use L-serine as substrate and share the identical gene ID. Based on the literature(Kulik et al., 2005), it can be confirmed that TRPS1 is an overall reaction catalyzed by whole tryptophan synthase, while TRPS2 is a partial reaction, which constitutes TRPS1 with the additional partial reaction TRPS3 (b1260 and b1261; 3ig3p_c --> g3p_c + indole_c) corresponding to the same genes in the model. Therefore, reaction TRPS1 should be retained, while the partial reactions TRPS3 and TRPS2, respectively catalyzed by the *α* and *β* subunits(David L. Nelson and Cox, 2008), should be closed.

After this adjustment, TRPAS2_reverse will be silenced by the shutdown of indole generation reaction TRPS3. According to the literature, the indole produced by the TRPS3 reaction will be transmitted to TRPS2 through a channel inside the enzyme (Kulik *et al*., 2005), so that TRPAS2_reverse cannot normally capture the substrate indole. When driven by a direct supply of highly concentrated ammonia, TRPAS2_reverse can be used for L-tryptophan synthesis(Watanabe and Snell, 1972), but its major function is L-tryptophan decomposition(Do et al., 2014). Furthermore, considering that the transmission of intermediate metabolites inside the enzyme molecule has overwhelming natural advantages, TRPAS2_reverse is inevitably eclipsed by TRPS1(Miles, 2001).

In addition, the MTRPOX reaction does not have the ability to synthesize L-tryptophan in the model, and its substrate *N*-methyltryptophan (nmtrp_c) only participates in this one reaction, that is, it can only be consumed and not regenerated, so that the reaction does not participate in the FBA calculation process in most cases (except when the substrate *N*-methyltryptophan is directly supplied). Accordingly, it is redundant in the model and is also not a reasonable candidate for L-tryptophan synthesis.

Returning to the analysis of MDF levels, since all the bottleneck reactions at stages 9 to 10 are central metabolic reactions, and the L-tryptophan synthesis process is abnormal due to improper expression of stoichiometric equations, the thermodynamic MDF levels of L-tryptophan pathway(s) should be redrawn after adjusting the stoichiometric framework based on the above analysis. The adjusted reactions are listed in table 2.

**Table 2.** Revised reactions in L-Trp synthesis process analysis

| Reaction ID | Reaction equation | Gene ID | Notes | $\Delta_r G_i'$<br>kJ/mol | $k_{cat}/MW$<br>h<sup>-1</sup>/kDa |
| --- | --- | --- | --- | --- | --- |
| TRPS1 | 3ig3p_c + ser__L_c --> g3p_c + h2o_c + trp__L_c | b1260 & | Keep | -30.5±6.2 | 145.44 |
| TRPS2 | indole_c + ser__L_c --> h2o_c + trp__L_c | b1261 | Shut | / | / |
| TRPS3 | 3ig3p_c --> g3p_c + indole_c |  | Shut | / | / |
| ACONTa | cit_c --> acon_C_c + h2o_c | b0118 & | Shut | / | / |
| ACONTb | acon_C_c + h2o_c --> icit_c | b1276 | Shut | / | / |
| ACONT | cit_c --> icit_c |  | Add | 8.3±0.6 | 3540 |
| ACONTa_r | acon_C_c + h2o_c --> cit_c |  | Shut | / | / |
| ACONTb_r | icit_c --> acon_C_c + h2o_c |  | Shut | / | / |
| ACONT_r | icit_c --> cit_c |  | Add | -8.3±0.6 | 3540 |

The combination of the two groups of reactions is shown in table 2, in which the reactions group of ACONT is reversible. The MDF information of the corrected L-tryptophan synthesis process is shown in table S11. Overall, while both the number of steps and the MDF values increased, the maximal flux decreases from 4.13 to 3.27 g/gDW/h, representing an obvious yield reduction as shown in Fig. 3.

Kishore et al. published a quantitative and systematic study of the apparent thermodynamic difference between the overall biochemical reaction and the separate reactions(Kishore et al., 1998) in the L-Trp synthesis process. To test the rationale of reaction adjustment, we visualized the L-tryptophan synthesis pathway as shown in Fig. 4. The pathway structure is consistent with both the work of Kishore et al.(Kishore *et al*., 1998) and the *E. coli* L-tryptophan synthesis pathway in the MetaCyc database. The synthesis pathway of L-tryptophan has three main confluence points, respectively corresponding to the ANS, ANPRT and TRPS1 reactions. For easy comparison, the pathways predicted by the *i*ML1515 model and the original EcoETM are shown in Figures S4 and S5, respectively. It can be seen that both pathways mistakenly use pyruvate and ammonia as direct substrates rather than L-serine. Therefore, although the introduction of enzymatic and thermodynamic constraints can reduce the yield solution space and reproduce the acetate overflow metabolism, the model is still not sufficiently realistic to avoid the prediction error of pathway structure and targets.

**Fig. 4.**
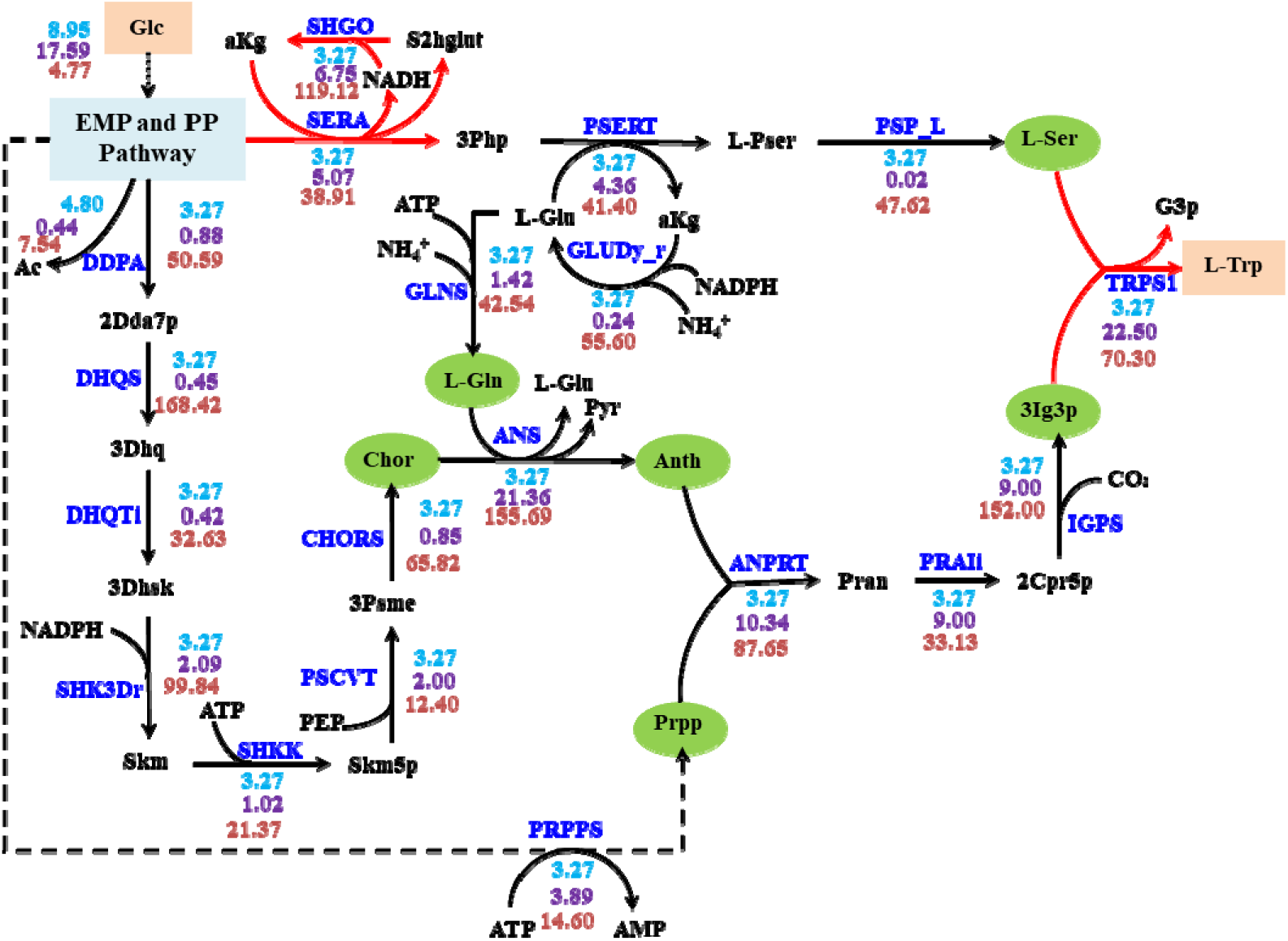
Predicted pathway of L-Trp synthesis. The revised reactions are indicated by red arrows, and the metabolites at the end of sub-pathway branches are indicated by a green background. The unit of the flux value is mmol/gDW/h (blue, on top), the unit of the enzyme cost is mg/gDW (purple, in the middle), and the unit of the maximal thermodynamic driving force is kJ/mol (orange, at the bottom). The upper limit of the glucose uptake rate was 10 mmol/gDW/h.

### 3.4 Analysis of the anthranilate synthesis process

The synthesis reaction of anthranilate (2-aminobenzoate, ANS) attracted our attention due to its high enzyme cost and glutamine consumption. Anthranilate is an important precursor of L-tryptophan, and its synthesis process is catalyzed by anthranilate synthase, which is widely present in microorganisms, including *E. coli*(Creighton and Yanofsky, 1970). It is reported that the enzyme can use both glutamine and ammonia as substrates(Tamir and Srinivasan, 1970) and is allosterically inhibited by L-tryptophan(Morollo and Bauerle, 1993). In 2001, Morollo and Eck mapped the mechanism of anthranilate synthase(Morollo and Eck, 2001). Ammonia is released by glutamine hydrolysis and transported through a channel inside the enzyme to trigger the ADCSN reaction. Subsequently, the intermediate metabolite amino-deoxy-isochorismate (ADIC) can be cleaved to produce pyruvate and anthranilate, as shown in Fig. 5A. Therefore, we selected the anthranilate synthesis process with a clear reaction mechanism to discuss the thermodynamic difference between glutamine and inorganic ammonia as substrates.

**Fig. 5.**
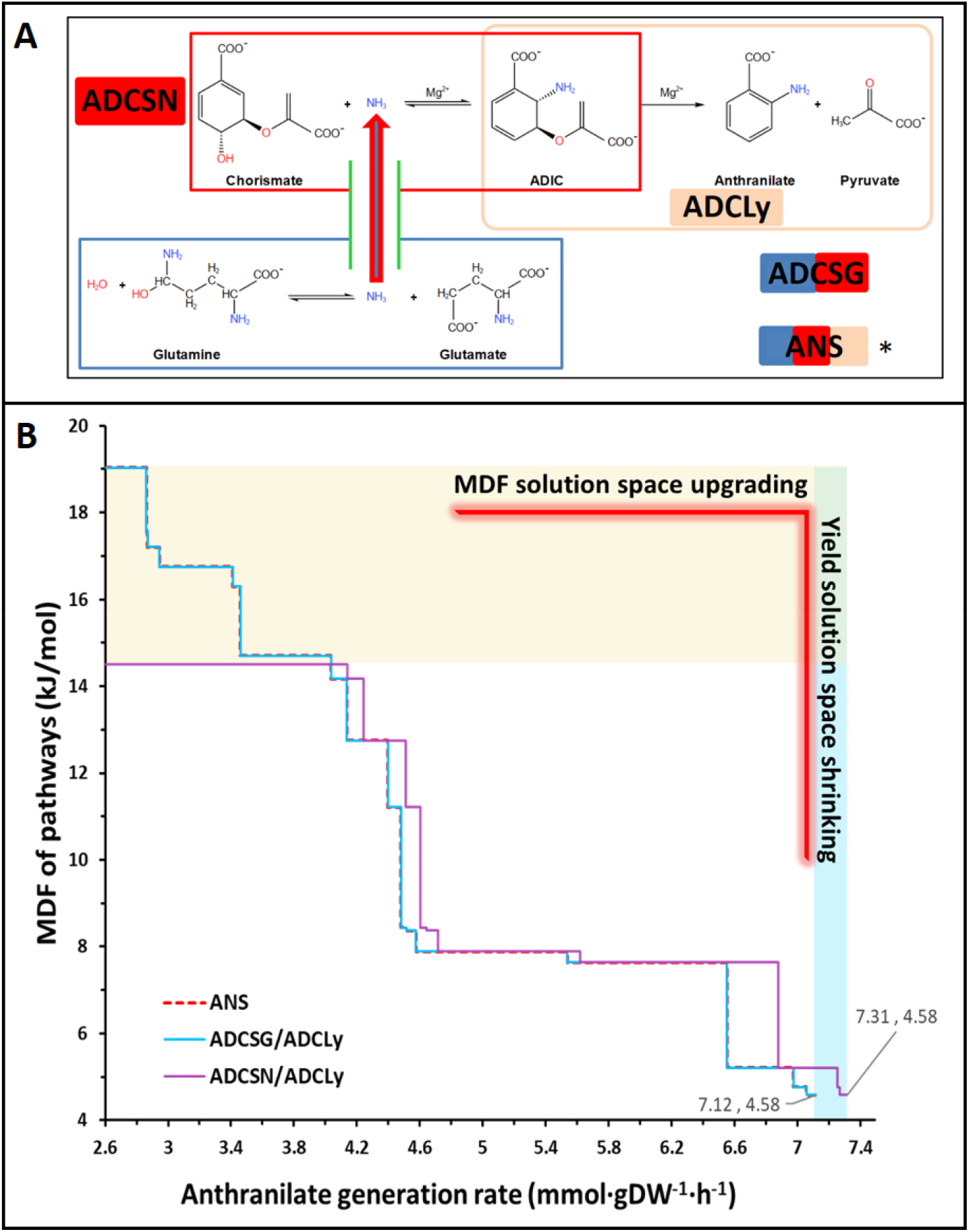
The synthetic process of anthranilate. **A**. The reaction mechanism of anthranilate synthase. The reaction with an asterisk (*) is the merged overall reaction in the *i*ML1515 model. The biological phenomena and principles reproduced in this figure refer to the work performed by Morollo and Eck(Morollo and Eck, 2001). **B**. MDF levels of different anthranilate synthesis processes. The purple curve shows the synthesis process with ammonia as substrate, while the red dotted line and blue line respectively indicate the two synthesis processes using glutamine as substrate (ANS is the combined reaction; ADCSG/ADCLy represent two partial reactions, and the curves show the undifferentiated result of these two processes).

In the *E. coli i*ML1515 model, anthranilate synthesis is achieved by a combined reaction (ANS). At the same time, there is a very similar synthesis process of 4-aminobenzoate in the model, which is represented by the two partial reactions ADCs and ACDL. Although the structures of 2-aminobenzoate and 4-aminobenzoate are very similar, their functions are quite different, since the latter mainly flows to the synthesis pathway of tetrahydrofolate (THFA). To compare the effects of glutamine and ammonia on the synthesis process of anthranilate, the partial reactions of ANS, including ADCSN, ADCSG and ADCLy, were added into the model (Fig. 5A) to simulate the process of direct utilization of ammonia and indirect utilization of glutamine. Accordingly, we added an intermediate metabolite ID, 2adcho_c (2-amino-4-deoxychorismate, KEGG No. c18054), as shown in tables S12 and S13.

According to the reaction combinations, anthranilate can be produced by the combined reaction ANS, or by two semi-reactions, in which the partial reaction to provide ammonia can be either ADCSG or ADCSN. Therefore, there can be three combinations: ①ANS; ②ADCSG and ADCLy, and ③ADCSN and ADCLy. Here, the corresponding Δ_*r*_*G*_*i*_′□ value was added for each reaction (table S12). ADCLy is a common and necessary reaction, so its kinetic parameters are set to be the same as the combined reaction ANS, while for the specific partial reactions ADCSG and ADCSN, the parameters were not set repeatedly, as shown in table S13.

When inorganic ammonia is used (③the combination of ADCSN and ADCLy), the pathway is thermodynamically favorable, and the anthranilate yield is higher than that of other two pathways with glutamine as substrate (were ①ANS; ②the reaction combination of ADCSG and ADCLy). In the two cases with glutamine as the substrate, both when the whole reaction and when the combination of two partial reactions is adopted, there is no difference of MDF at each yield level (Fig. 5B), since the partial reaction does not produce new pathway combination(s). At the same expected flux level, the MDF of the two pathways with glutamine as substrate always decreases earlier than with ammonia as substrate, and the flux solution space is reduced, as shown in table S14. It can be preliminarily concluded that the process using ammonia as substrate is thermodynamically feasible and relatively economical in terms of carbon yield, as shown in Fig. 5B. In the two pathways with glutamine as substrate, there are five stages in which the MDF level is better than that of the pathway with ammonia as substrate in the low yield stages. The reason is that the optimal thermodynamic driving force of ASCDN is 14.514 kJ/mol, which has become the MDF ceiling of the pathway(s) in which it participates (Fig. 5B and table S14).

However, such an advantage does not seem to be a necessity for survival. Therefore, it is more convincing to understand the significance of glutamine indirectly providing ammonium from the perspective of avoiding the toxicity of inorganic ammonium, increasing the affinity of the enzyme for the amino group and increasing the storage capacity of nitrogen sources as an adaptation to nitrogen starvation and acute acid stress(Lu et al., 2013; van Heeswijk et al., 2013). However, it is unreasonable to assume that the high-toxicity, low-affinity and unstable supply of ammonium will not lead to the poor thermodynamic feasibility of the cell growth process. We believe that reliable application calls for the deepening of the constraint integration principle and the refinement of various relevant parameters in the reconstruction process of constraint-based GEMs(Bi et al., 2022). For example, the abiotic constraints (ABCs) model(Akbari et al., 2021) was applied as a constraints-refined framework to the core model of *E. coli*, albeit currently consisting of only 78 reactions. Predictably, constraint-based models still have a long way to go in terms of accuracy improvement, optimization of principles and scale magnification.

## 4. Discussion

Incomplete reactions in GEMs lead to unrealistic opportunities to participate in the pathway alone and increase the number of reaction combinations, which will expand the solution space of synthetic pathways. However, the separation of linked reactions will cause the exposure of intermediate metabolites and strengthen the constraints, which is unfavorable to the solution space of thermodynamic feasibility(Noor *et al*., 2014). Conversely, when the partial reactions are combined, the pathway yield may be partially lost due to limitations of thermodynamic feasibility. The synthesis examples of L-serine and L-tryptophan illustrate the trade-off between pathway yield and thermodynamic driving force.

In previous studies on thermodynamics, due to the lack of Gibbs free energy parameters of some reactions, the exposure of intermediate metabolites with yet unquantifiable energy level was reduced by combining reactions(Kiparissides and Hatzimanikatis, 2017; Noor et al., 2013). Similarly, during the determination of enzyme kinetic parameters, the exposure of undetectable intermediate metabolites can also be avoided by using more easily detected and more stable end products as indicators by coupling reactions(Corbalán-García et al., 1993; Yang *et al*., 2019). In the process of network analysis, the combination of reactions can also avoid the problem of combinatorial explosion, making it computationally feasible to analyze the main network topology(Liu and Sumpter, 2018). Thus, combined chemical reactions can provide feasibility and convenience for the study of several problems. In this work, the same small adjustment of combining reactions helped us compare overall reactions and partial reactions.

As shown in Fig. 3, the trade-off between yield and MDF has two layers of meaning, since it 1) exists inside each curve, and also 2) exists between two curves. The fundamental reason is identical for both the combination of reactions and the setting of high yield essentially excluding the possibility of some reaction combinations (i.e., pathways). Accordingly, both have the function of reducing the solution space. However, the combination of reactions is beneficial to the improvement of the thermodynamic driving force and unfavorable to the maintenance of the yield. Conversely, a high yield level setting is beneficial to maintain the yield, but unfavorable to maintain the thermodynamic driving force level.

Multifunctional enzymes or enzyme complexes can also be considered as cell compartmentalization strategies, which is of great significance for the feasibility of metabolic processes. This concept should be fully referenced in both the construction and application of constraints-based GEMs. At present, the enzyme-constrained models only consider the effects of multifunctional enzymes and enzyme complexes in terms of the molecular weight of proteins(Adadi *et al*., 2012; Sanchez *et al*., 2017), but underestimate the biologically meaningfully evolved efficiency of complex enzyme molecules such as pyruvate dehydrogenase (PDH) and 2-ketoglutarate dehydrogenase (AKGDH)(Mao *et al*., 2022), resulting in the overestimation of enzyme costs, which distorts the predicted pathway structure (the real optimal pathway is replaced by the suboptimal pathway due to the estimated deviation of enzyme cost) and/or flux (even if there is no false switching of pathways, the flux will be reduced due to excessive resource constraints), and makes pathway evaluation(Desouki et al., 2015; Du et al., 2018a) unreliable.

In addition, if the microscopic compartmentalization concept is reasonably adopted, more engineering possibilities may also be realized. For example, the enzyme complex encoded by the *fad*AB genes in the fatty acid *β*-oxidation pathway(Dellomonaco et al., 2011) can be disassembled, so that the intermediate metabolite *β*-ketoacyl-CoA can be released and then integrated into a variety of bio-polyesters(Olivera et al., 2001). In addition, the internal structure of the enzyme can not only enable compartmentalization, but the undisturbed microenvironment can provide conditions for the local accumulation of metabolites (Durrani et al., 2020; Srere and Mosbach, 1974). Therefore, although there is no structural coupling relationship between some enzyme functions such as internal channels, there may nevertheless promote the thermodynamic feasibility, such as cascade reactions catalyzed by the “fused diaminohydroxyphosphoribosylaminopyrimidine deaminase and 5-amino-6-(5-phosphoribosylamino)uracil reductase” (Magalhães et al., 2008) and “fused N-acetylglucosamine-1-phosphate uridyltransferase and glucosamine-1-phosphate acetyltransferase” (Gehring et al., 1996) expressed by the genes b0414 and b3730, respectively. Thus, it is important to set the parameters in the process of model correction and analysis based on the real intracellular conditions to obtain effective prediction information. At the same time, it is meaningful to investigate the structure and catalytic mechanism of enzymes(Crampin et al., 2004; Koenig et al., 1961), as typical molecular machines(Demirel, 2010; Elber and Kirmizialtin, 2013), to help us understand the intermediate metabolite(s) that mediate the relationship between reactions from multiple aspects. These include, 1) whether they will separate from enzyme molecule, 2) whether they will escape from the enzyme structure and the escape efficiency, as well as 3), whether the environment facilitates their diffusion, which will provide an important basis for us to estimate the degree of thermodynamic promotion between reactions. Accordingly, improved in-depth analysis of protein structure and function is crucial for the simulation and prediction of cell behaviors(Ornes, 2022).

In addition, the bottleneck reactions predicted in this study often include reactions in the central metabolic pathways, which may help us understand why there are oscillations in central metabolic processes such as glycolysis(Gustavsson et al., 2012; Maria, 2020; Özsezen et al., 2019). All biochemical reactions in organisms cannot occur simultaneously due to constraints of thermodynamic feasibility and resource availability, just as all trains in a country cannot run simultaneously. Therefore, oscillations provide overall planning and coordination for the inner workings of the cellular system. This seems to be contrary to the theoretical basis of GEMs, which are based on the steady-state hypothesis and flux balance analysis(Orth et al., 2010), but just as computers will not operate in the same way as the human brain, this difference can be understood and accepted, so that nonequilibrium theory and the steady-state hypothesis have been and will continue to coexist and guide our reasoning(Colombo and Palacios, 2021; Wang et al., 2008).

The intelligence of cells is beyond our imagination, and how to reproduce cell space-time in digital cells will also be an interesting challenge, whereby whole-cell modeling(Karr et al., 2012) arguably represents the most hopeful vision. In addition to compartmentalization, cells may also obtain additional energy at high temperature, which can enable processes that are infeasible at room temperature, such as in some archaeal and thermophilic microorganisms(Kawarabayasi et al., 1998). Perhaps we should be more optimistic in our assessment and verification of pathway feasibility(Petrovic et al., 2018).

## Conclusions

Integrating the availability of enzyme resources and thermodynamic feasibility should improve the prediction accuracy of constraint-based GEMs. However, based on the analysis of the prediction anomaly in the pathway structure of L-serine synthesis, was found that there is a lack of consideration of multifunctional enzymes and enzyme complexes as a strategy of cell compartmentalization in the constraint-based GEMs. Accordingly, the unreasonable leakage of intermediate metabolites and the resulting unreasonable constraints can be effectively avoided by combining some incomplete or partial reactions. Finally, the structure of the corresponding synthetic pathways, such as those of L-serine and L-tryptophan, was corrected. At the same time, ideas and suggestions are provided for the treatment of reactions catalyzed by multifunctional enzymes or enzyme complexes. Hence, this work will have a far-reaching influence on the improvement of the structure and application for constraint-based GEMs by provided quantifiable evidence and a workable analysis method.

## Acknowledgement

This work was funded by the National Key Research and Development Program of China (2018YFA0900300, 2020YFA0908301), the National Natural Science Foundation of China (32201188), the Tianjin Synthetic Biotechnology Innovation Capacity Improvement Project (TSBICIP-CXRC-060 and TSBICIP-PTJS-001).

## Competing interests

The authors declare no competing financial interests.

## Author contributions

HM and XY conceived the project. XY and ZM developed the ETGEMs and constructed the EcoETM model. XY performed the calculation and visualization. YZ, JH and HD contributed to the analysis and discussion. ZM and RW maintains the calculation tool. HM, XY and YZ wrote the manuscript. All authors read and approved the manuscript.

## Supplementary information

See Supplementary data.

## Notes

### Competing Interest Statement

The authors have declared no competing interest.

