## Appendix for "The necessity of considering enzymes as compartments in constraint-based genome-scale metabolic models"

**Supplementary material**

**Table S1** Thermodynamic bottleneck reaction(s) in the l-serine synthesis pathways

| **Turning** | **Max flux** | **MDF** | **Bottleneck** | **Reaction equation** |
| --- | --- | --- | --- | --- |
| **point** | **(mmol/gDW/h)** | **(kJ/mol)** | **reaction** |  |
| 1 | 5.681 | 9.864 | ACONTa | cit_c --> acon_C_c + h2o_c |
|  |  |  | ACONTb | acon_C_c + h2o_c --> icit_c |
| 2 | 5.916 | 9.358 | PGI_r | f6p_c --> g6p_c |
|  |  |  | RPE | ru5p__D_c --> xu5p__D_c |
|  |  |  | G6PDH2r | g6p_c + nadp_c --> 6pgl_c + h_c + nadph_c |
|  |  |  | TALA | g3p_c + s7p_c --> e4p_c + f6p_c |
|  |  |  | TKT2 | e4p_c + xu5p__D_c --> f6p_c + g3p_c |
|  |  |  | TKT1 | r5p_c + xu5p__D_c --> g3p_c + s7p_c |
|  |  |  | TPI_r | g3p_c --> dhap_c |
|  |  |  | FBA_r | dhap_c + g3p_c --> fdp_c |
| 3 | 5.923 | 7.638 | ADK1 | amp_c + atp_c --> 2.0 adp_c |
| 4 | 8.306 | 7.422 | PGK_r | 13dpg_c + adp_c --> 3pg_c + atp_c |
|  |  |  | PGM_r | 3pg_c --> 2pg_c |
|  |  |  | GAPD | g3p_c + nad_c + pi_c --> 13dpg_c + h_c + nadh_c |
|  |  |  | ENO | 2pg_c --> h2o_c + pep_c |
|  |  |  | ASPK | asp__L_c + atp_c --> 4pasp_c + adp_c |
|  |  |  | ASAD_r | 4pasp_c + h_c + nadph_c --> aspsa_c + nadp_c + pi_c |
| 5 | 8.390 | 5.208 | PGK_r | 13dpg_c + adp_c --> 3pg_c + atp_c |
|  |  |  | PGM_r | 3pg_c --> 2pg_c |
|  |  |  | GAPD | g3p_c + nad_c + pi_c --> 13dpg_c + h_c + nadh_c |
|  |  |  | ENO | 2pg_c --> h2o_c + pep_c |
|  |  |  | TPI | dhap_c --> g3p_c |
| 6 | 8.919 | 4.767 | PGK_r | 13dpg_c + adp_c --> 3pg_c + atp_c |
|  |  |  | PGM_r | 3pg_c --> 2pg_c |
|  |  |  | GAPD | g3p_c + nad_c + pi_c --> 13dpg_c + h_c + nadh_c |
|  |  |  | ENO | 2pg_c --> h2o_c + pep_c |
|  |  |  | TPI | dhap_c --> g3p_c |
|  |  |  | FBA | fdp_c --> dhap_c + g3p_c |
| 7 | 8.994 | 4.584 | MDH | mal__L_c + nad_c --> h_c + nadh_c + oaa_c |
|  |  |  | FUM | fum_c + h2o_c --> mal__L_c |
| 8 | 17.175 | 1.567 | PGCD | 3pg_c + nad_c --> 3php_c + h_c + nadh_c |
| 9 | 20.079 | 0.522 | ENO_r | h2o_c + pep_c --> 2pg_c |
|  |  |  | PGM | 2pg_c --> 3pg_c |
|  |  |  | PGCD | 3pg_c + nad_c --> 3php_c + h_c + nadh_c |
| 10 | 20.119 | 0.219 | PGCD | 3pg_c + nad_c --> 3php_c + h_c + nadh_c |
|  |  |  | PGK_r | 13dpg_c + adp_c --> 3pg_c + atp_c |
|  |  |  | FLDR2 | 2.0 flxso_c + nadph_c --> 2.0 flxr_c + h_c + nadp_c |
|  |  |  | POR5_r | accoa_c + co2_c + 2.0 flxr_c + h_c --> coa_c + 2.0 flxso_c + pyr_c |
|  |  |  | ENO_r | h2o_c + pep_c --> 2pg_c |
|  |  |  | PPS | atp_c + h2o_c + pyr_c --> amp_c + 2.0 h_c + pep_c + pi_c |
|  |  |  | GAPD | g3p_c + nad_c + pi_c --> 13dpg_c + h_c + nadh_c |
|  |  |  | PGM | 2pg_c --> 3pg_c |
| 11 | 20.661 | 0.069 | PGK_r | 13dpg_c + adp_c --> 3pg_c + atp_c |
|  |  |  | TPI | dhap_c --> g3p_c |
|  |  |  | PGCD | 3pg_c + nad_c --> 3php_c + h_c + nadh_c |
|  |  |  | GAPD | g3p_c + nad_c + pi_c --> 13dpg_c + h_c + nadh_c |
| 12 | 20.833 | -0.482 | PGK_r | 13dpg_c + adp_c --> 3pg_c + atp_c |
|  |  |  | PGCD | 3pg_c + nad_c --> 3php_c + h_c + nadh_c |
|  |  |  | GAPD | g3p_c + nad_c + pi_c --> 13dpg_c + h_c + nadh_c |
|  |  |  | TPI | dhap_c --> g3p_c |
|  |  |  | FBA | fdp_c --> dhap_c + g3p_c |
| 13 | 21.029 | -0.632 | PFL | coa_c + pyr_c --> accoa_c + for_c |
|  |  |  | FLDR2 | 2.0 flxso_c + nadph_c --> 2.0 flxr_c + h_c + nadp_c |
|  |  |  | POR5_r | accoa_c + co2_c + 2.0 flxr_c + h_c --> coa_c + 2.0 flxso_c + pyr_c |

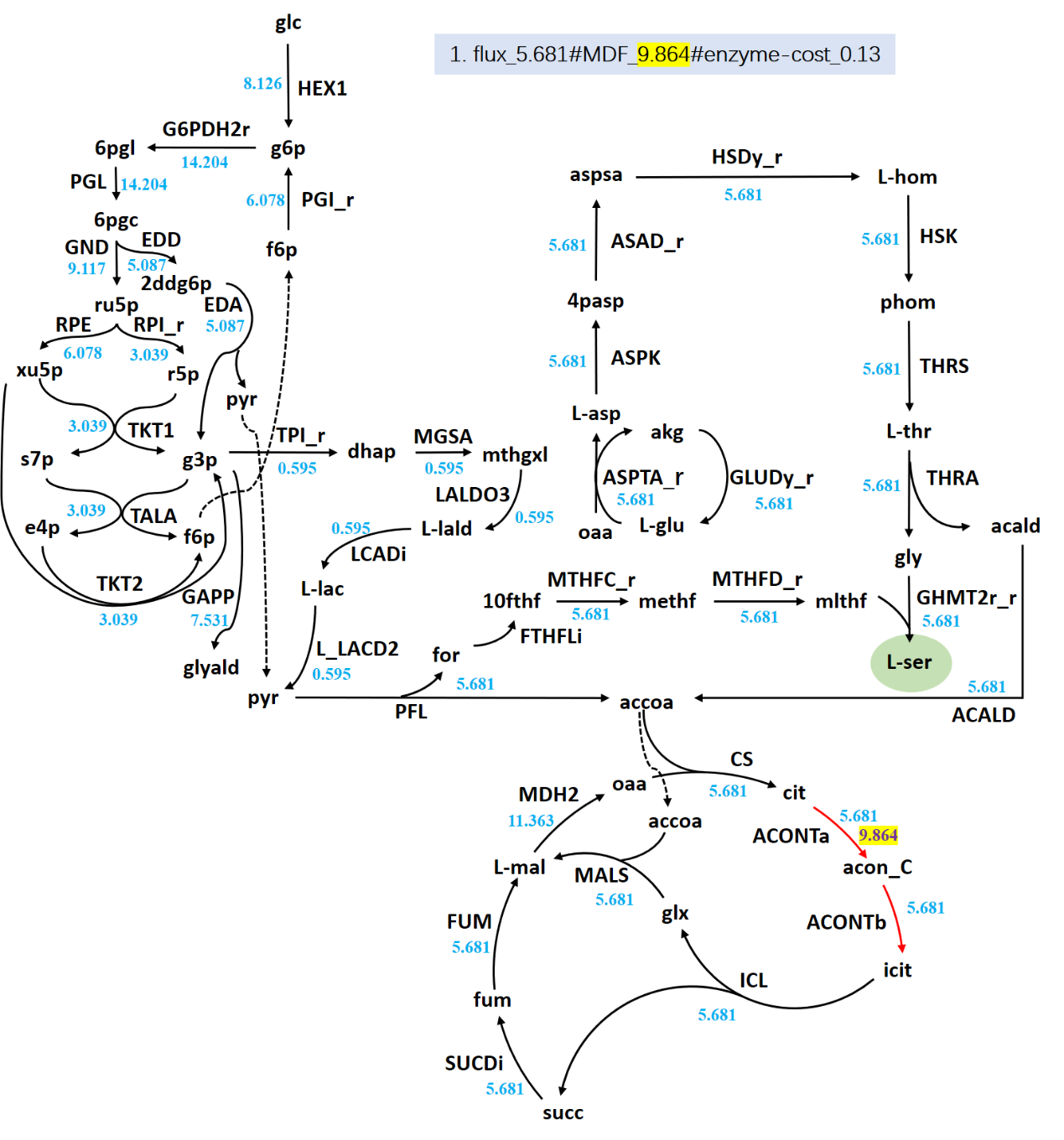

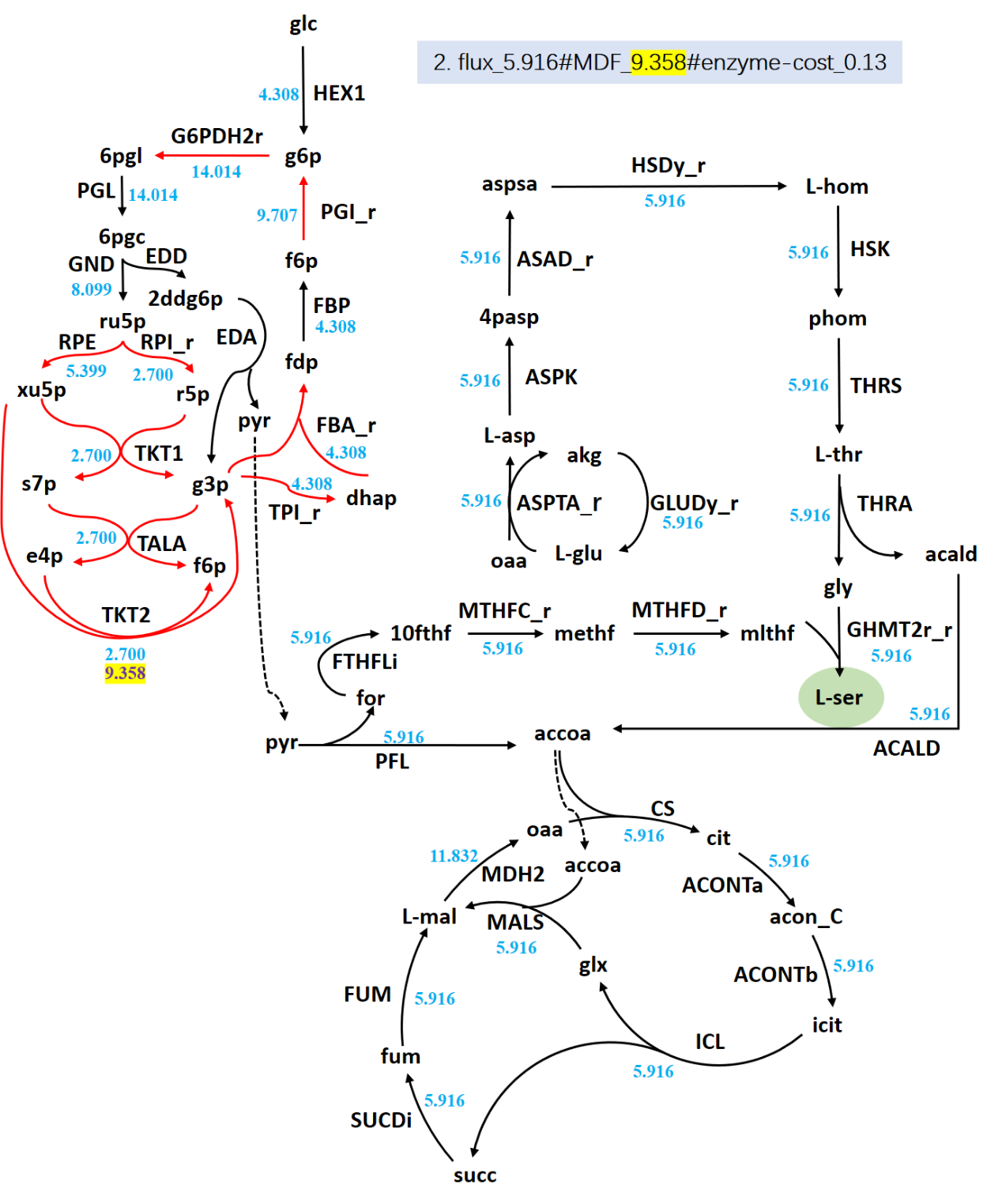

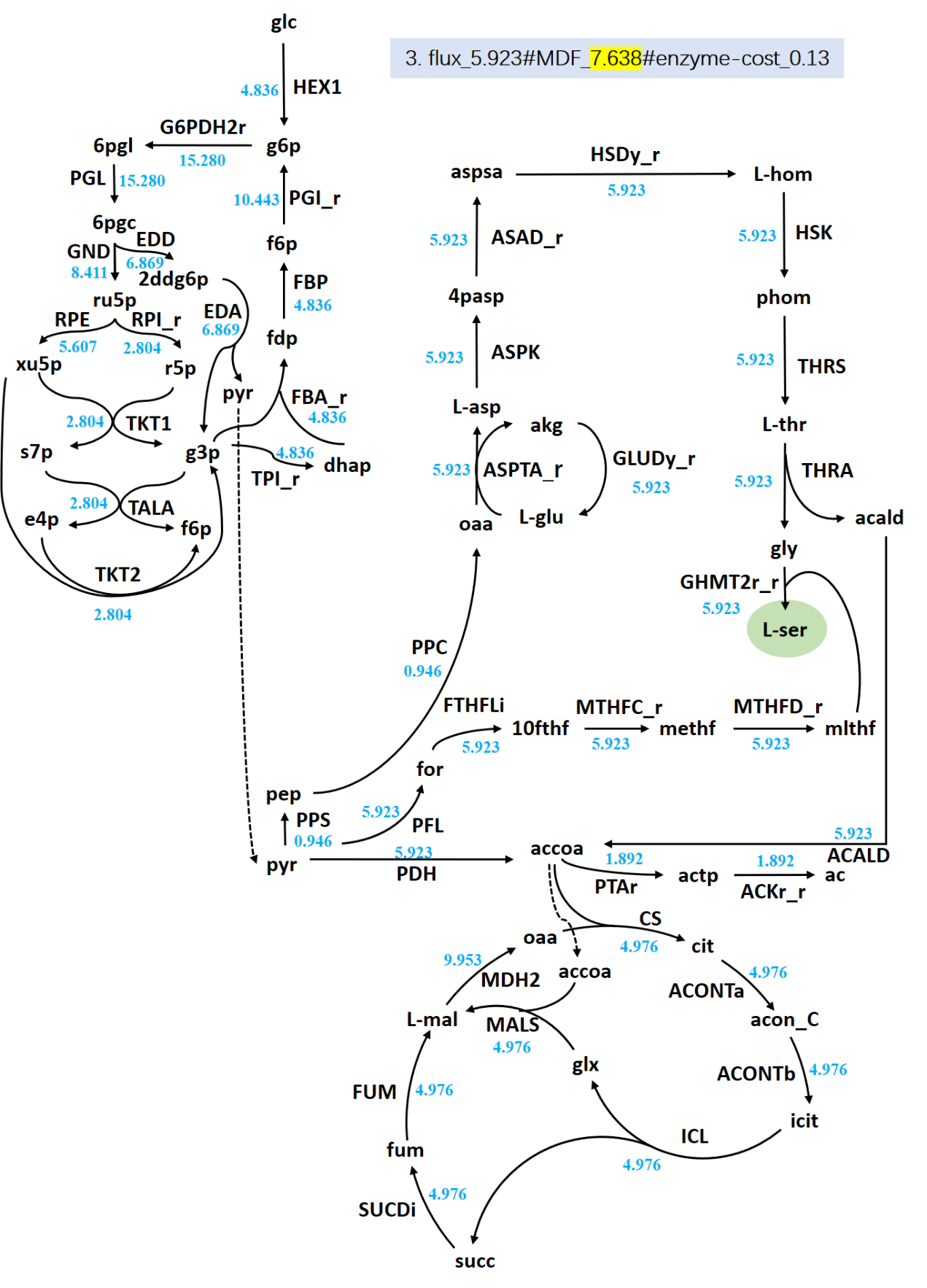

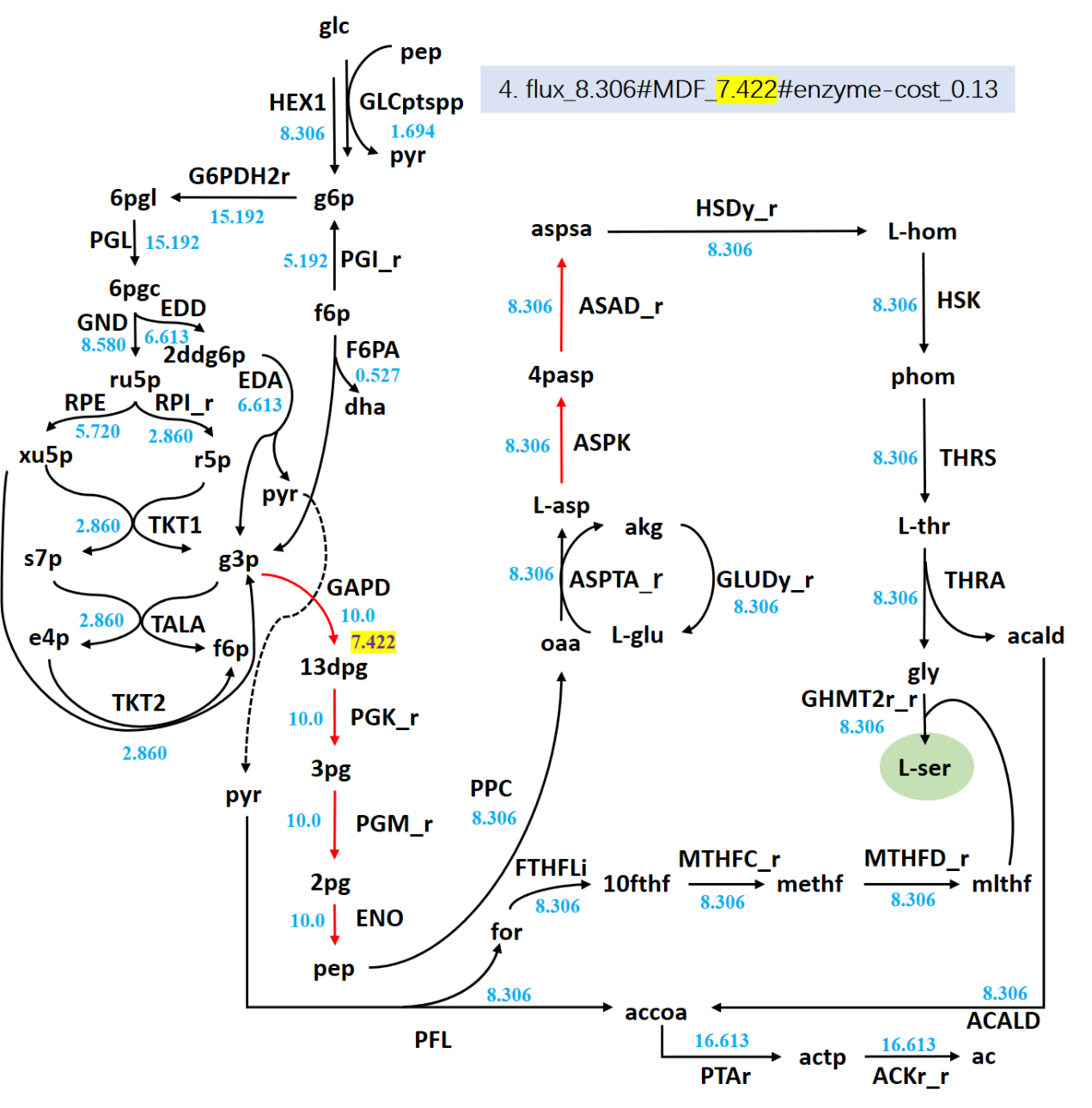

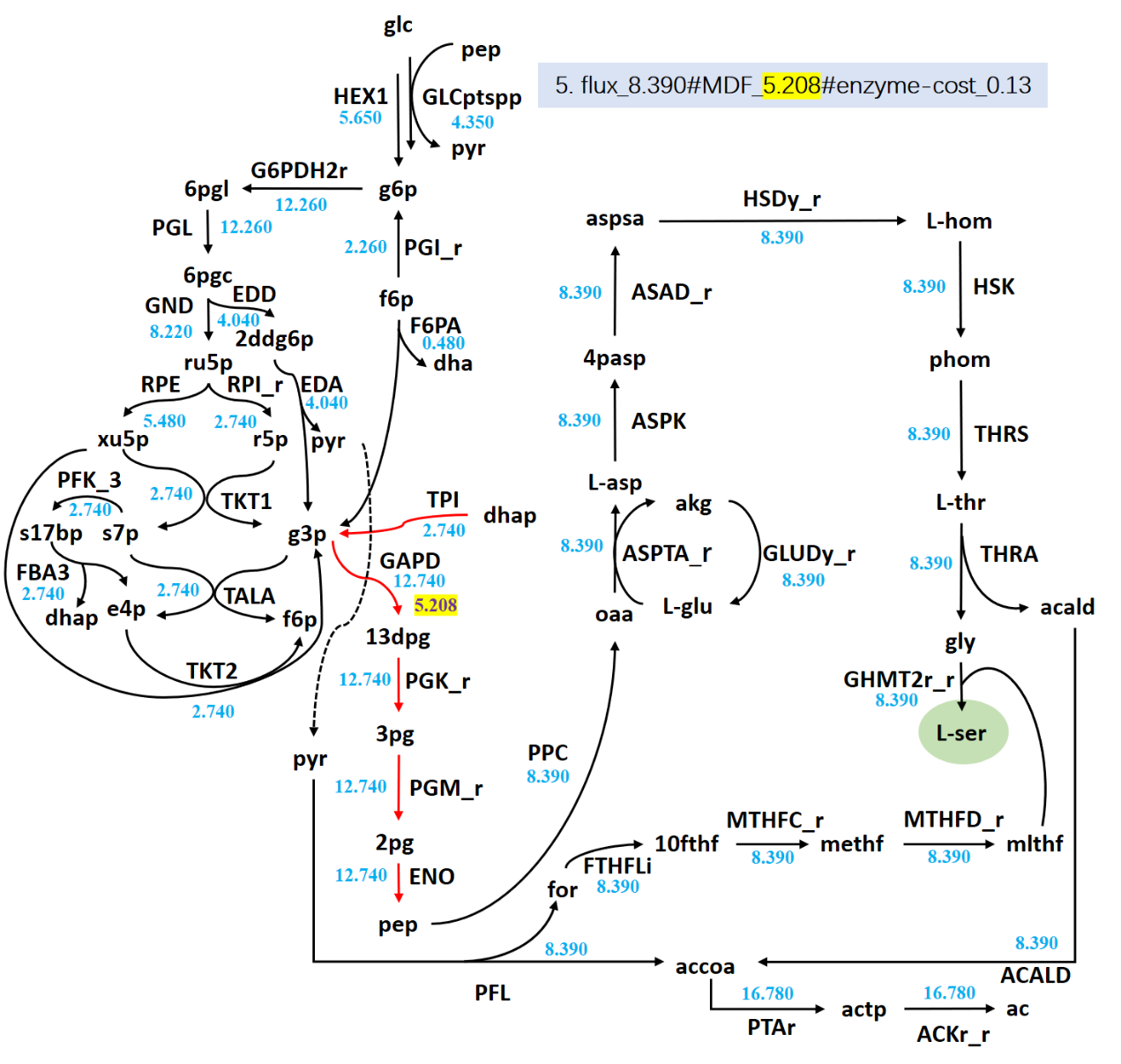

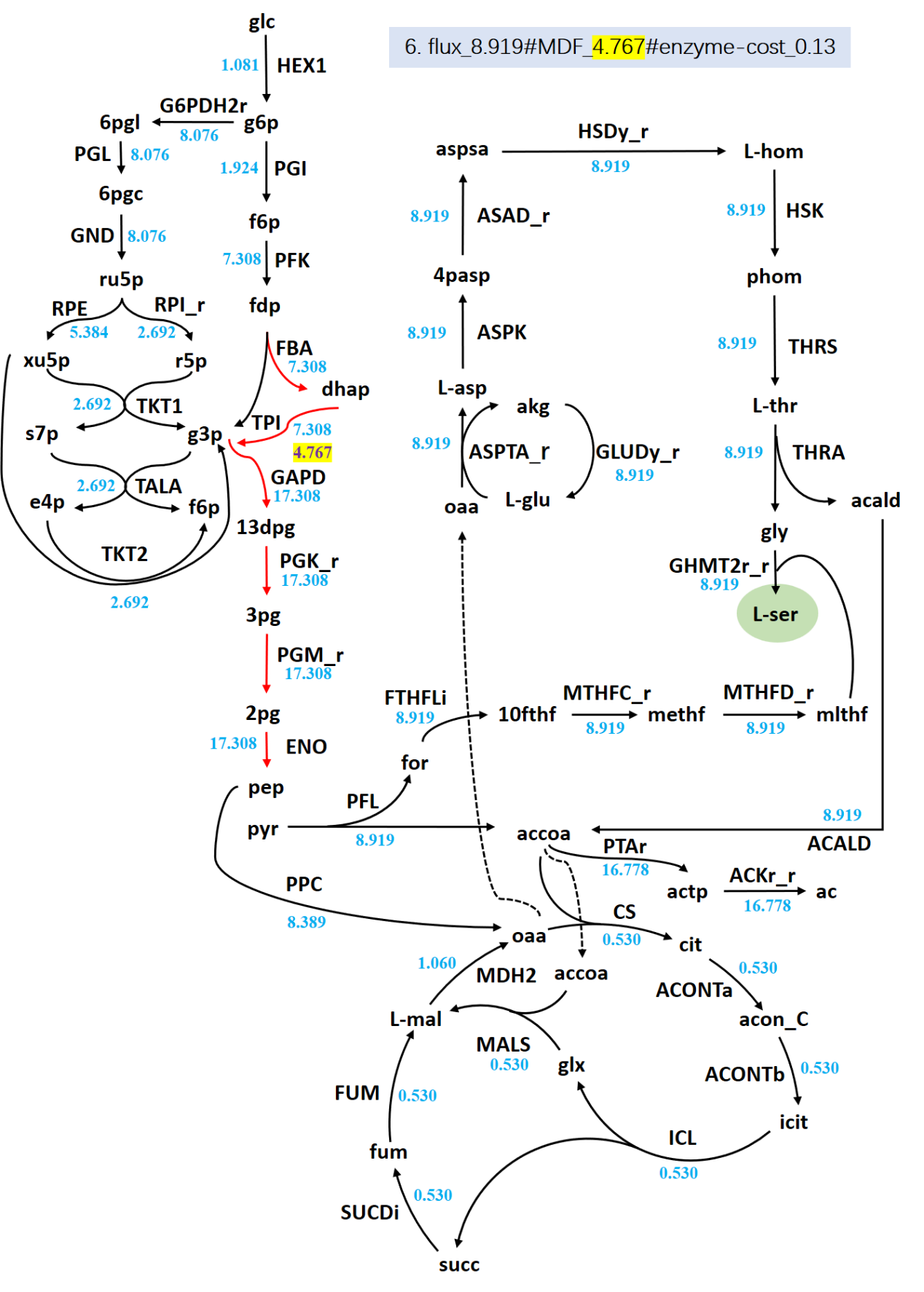

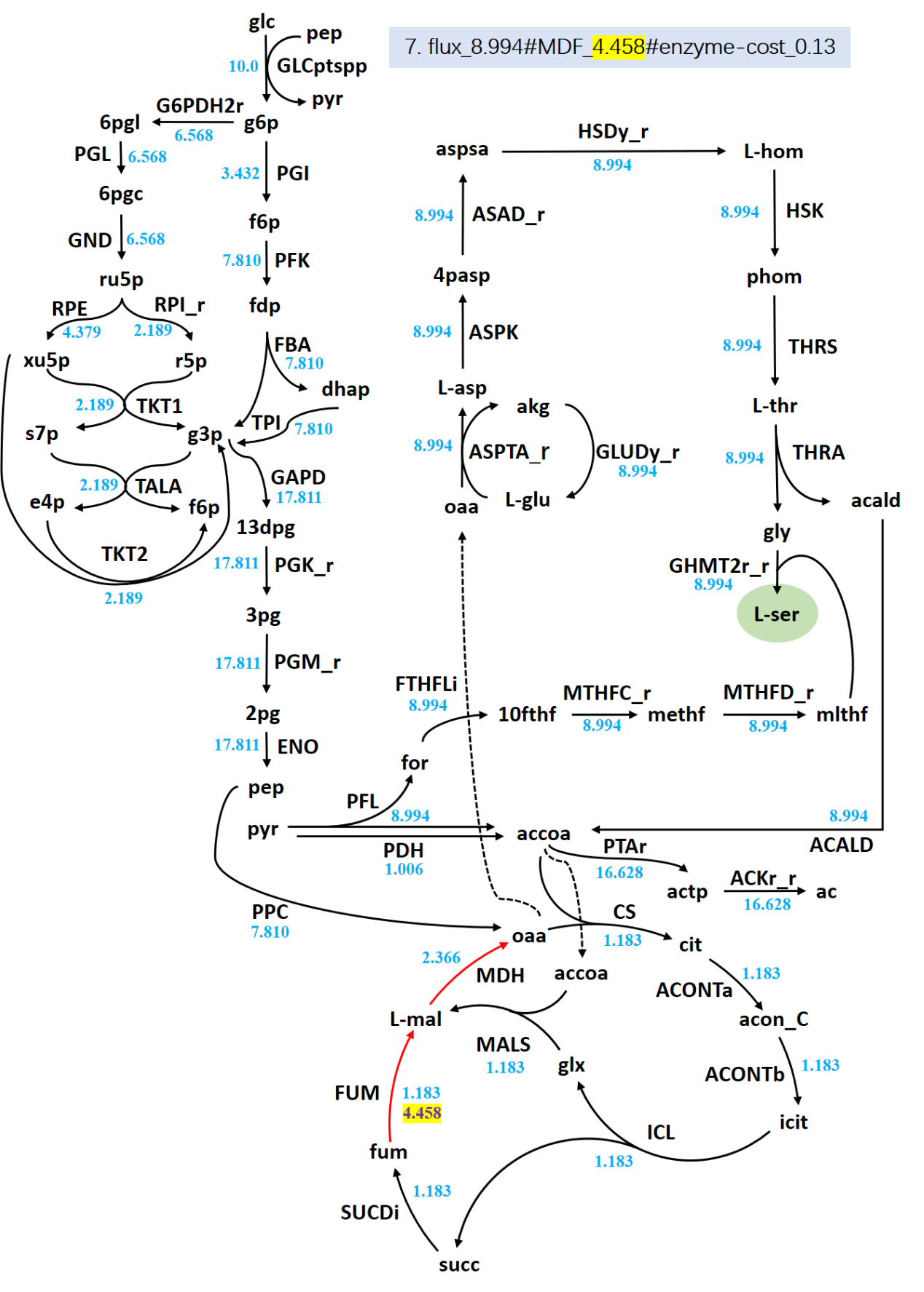

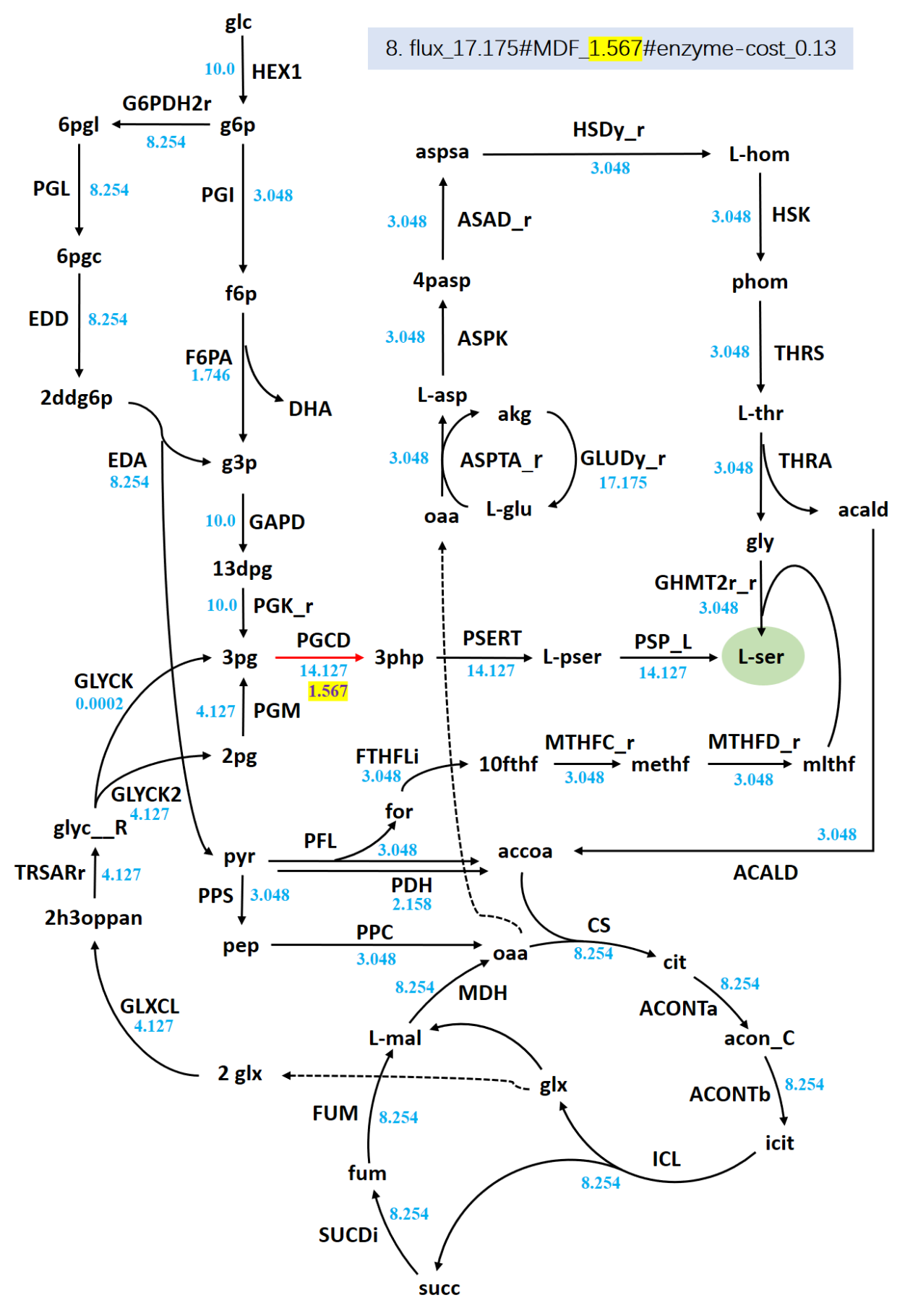

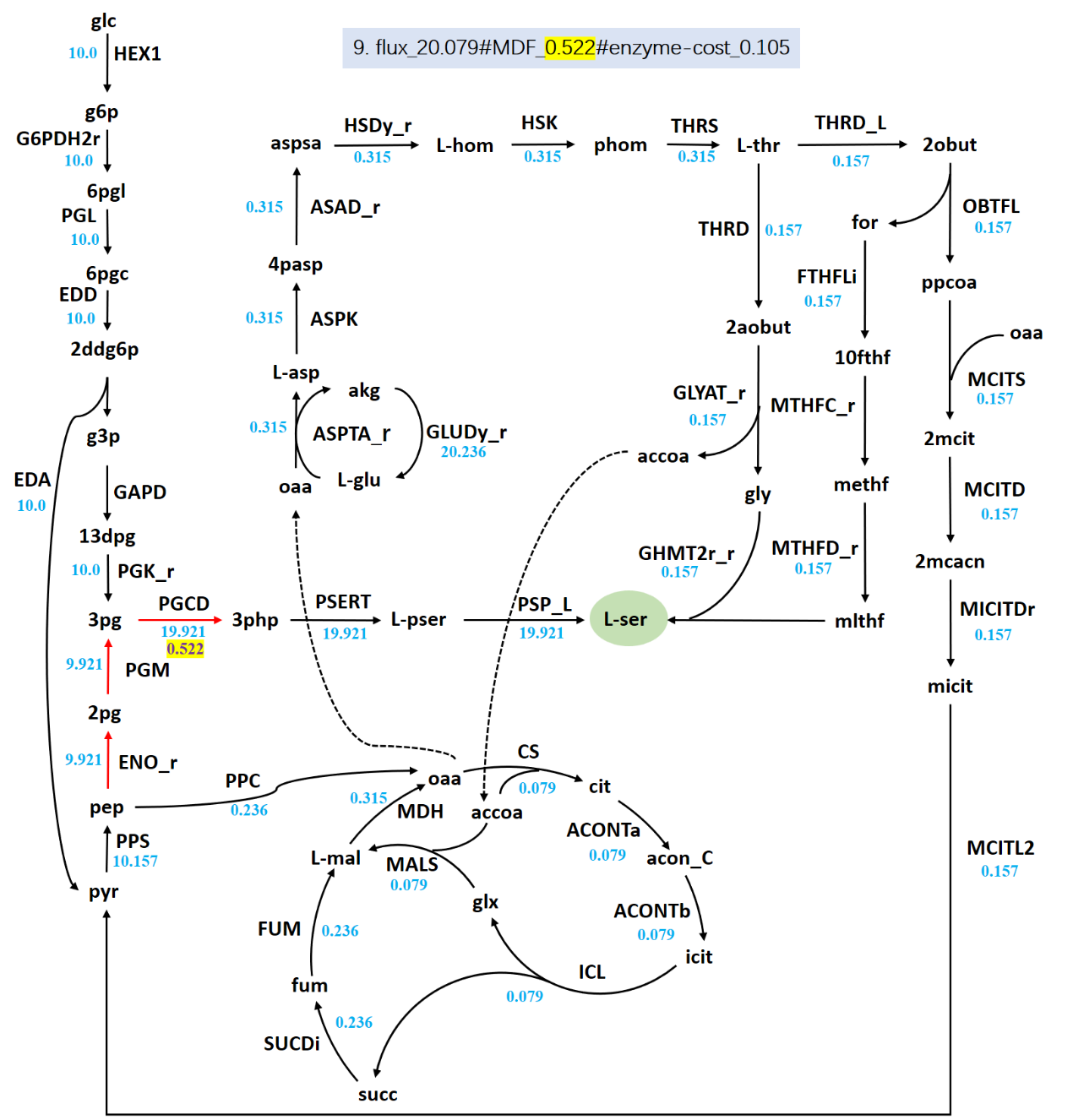

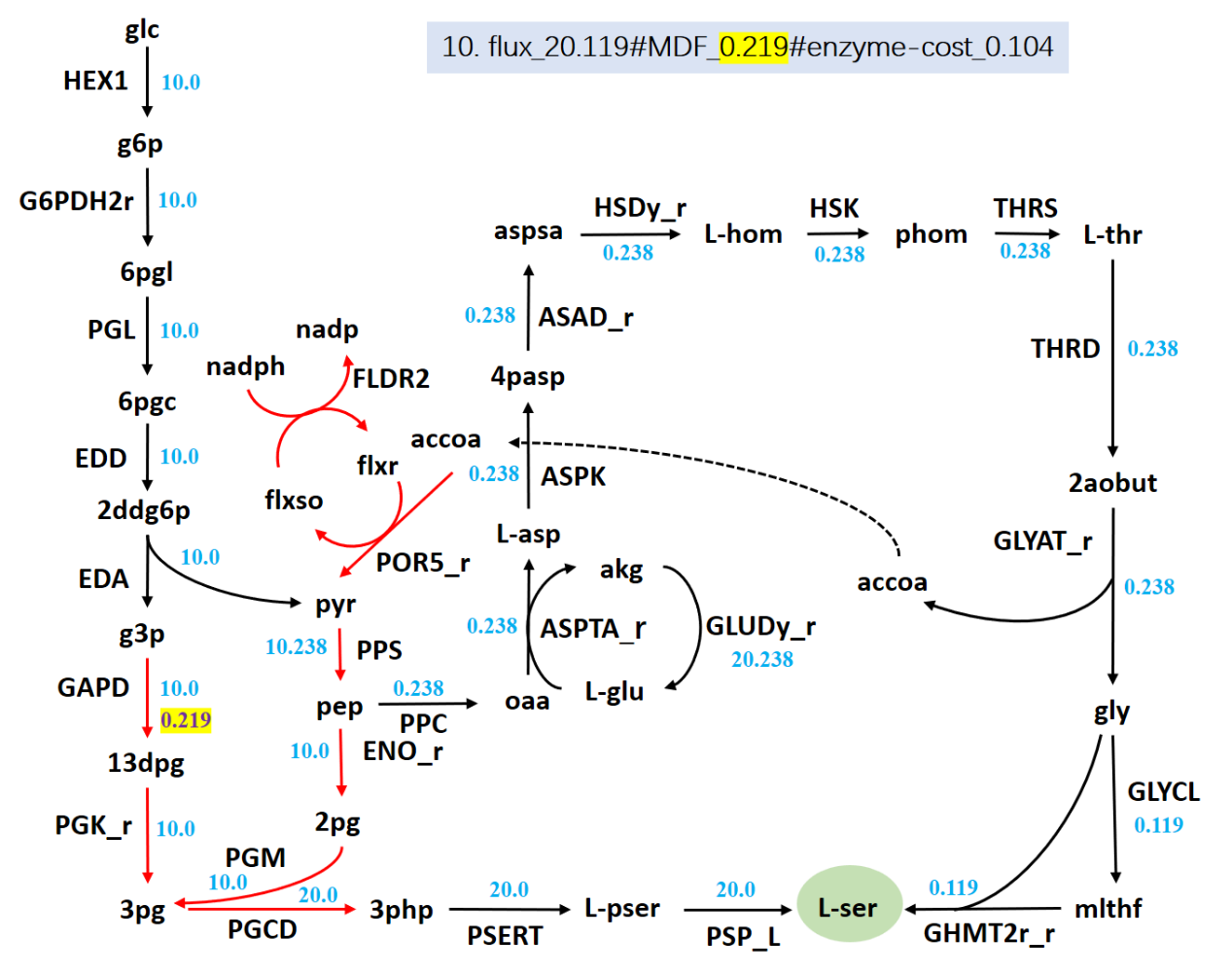

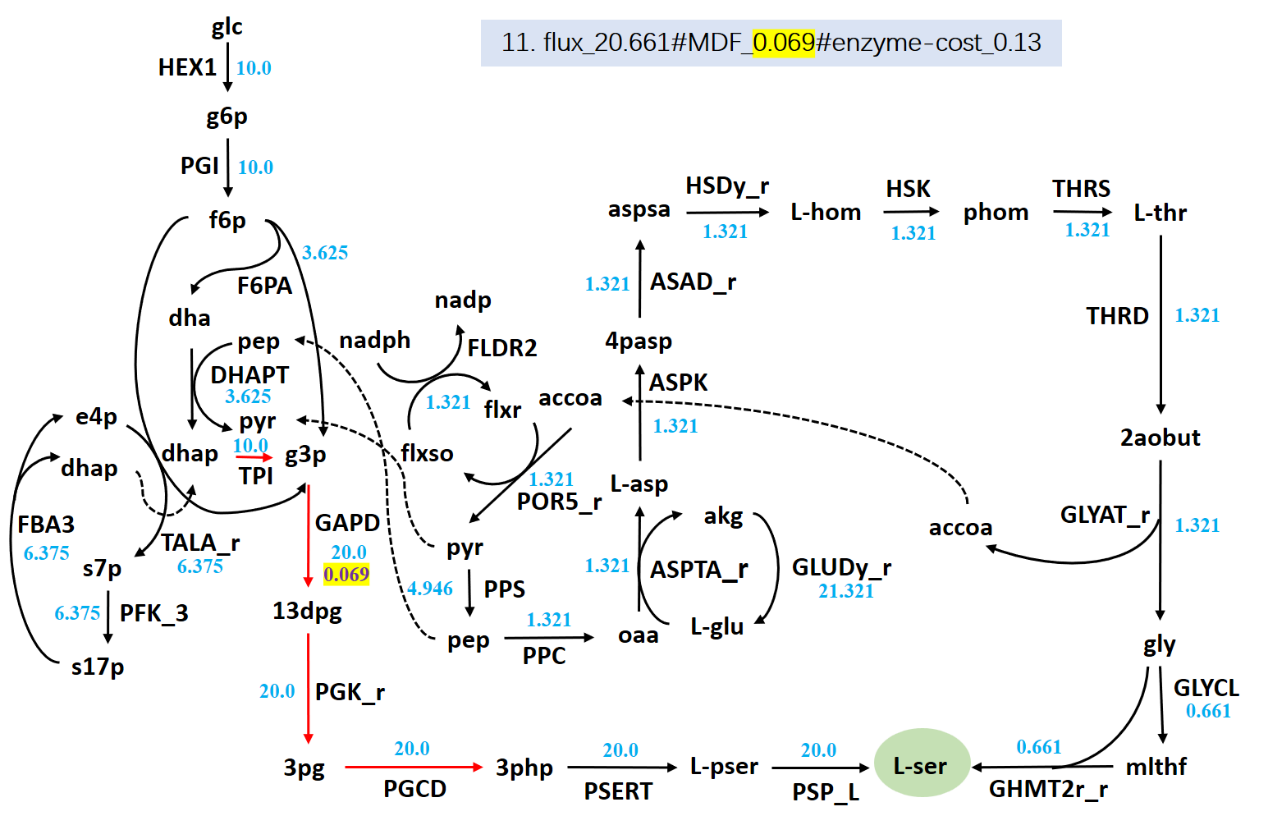

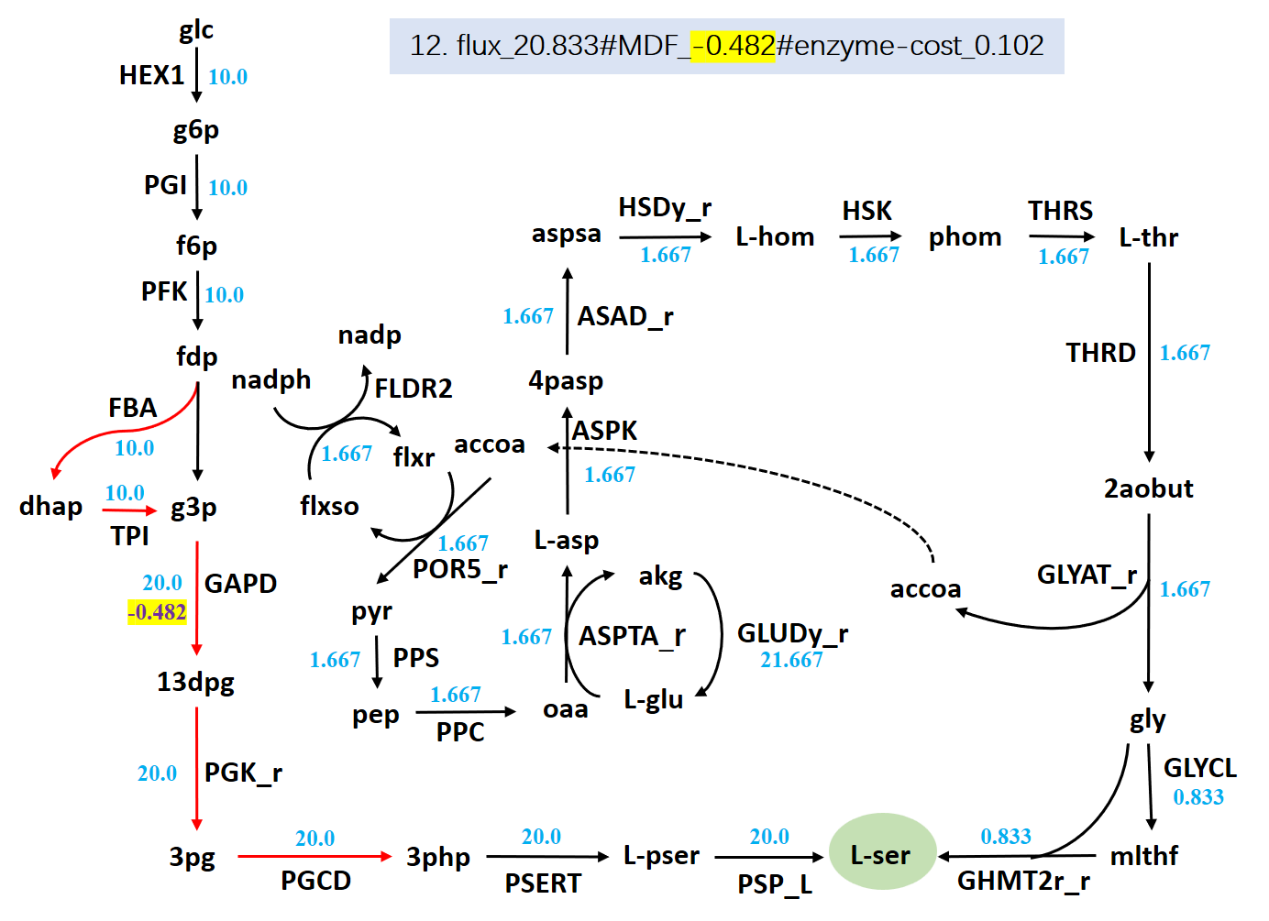

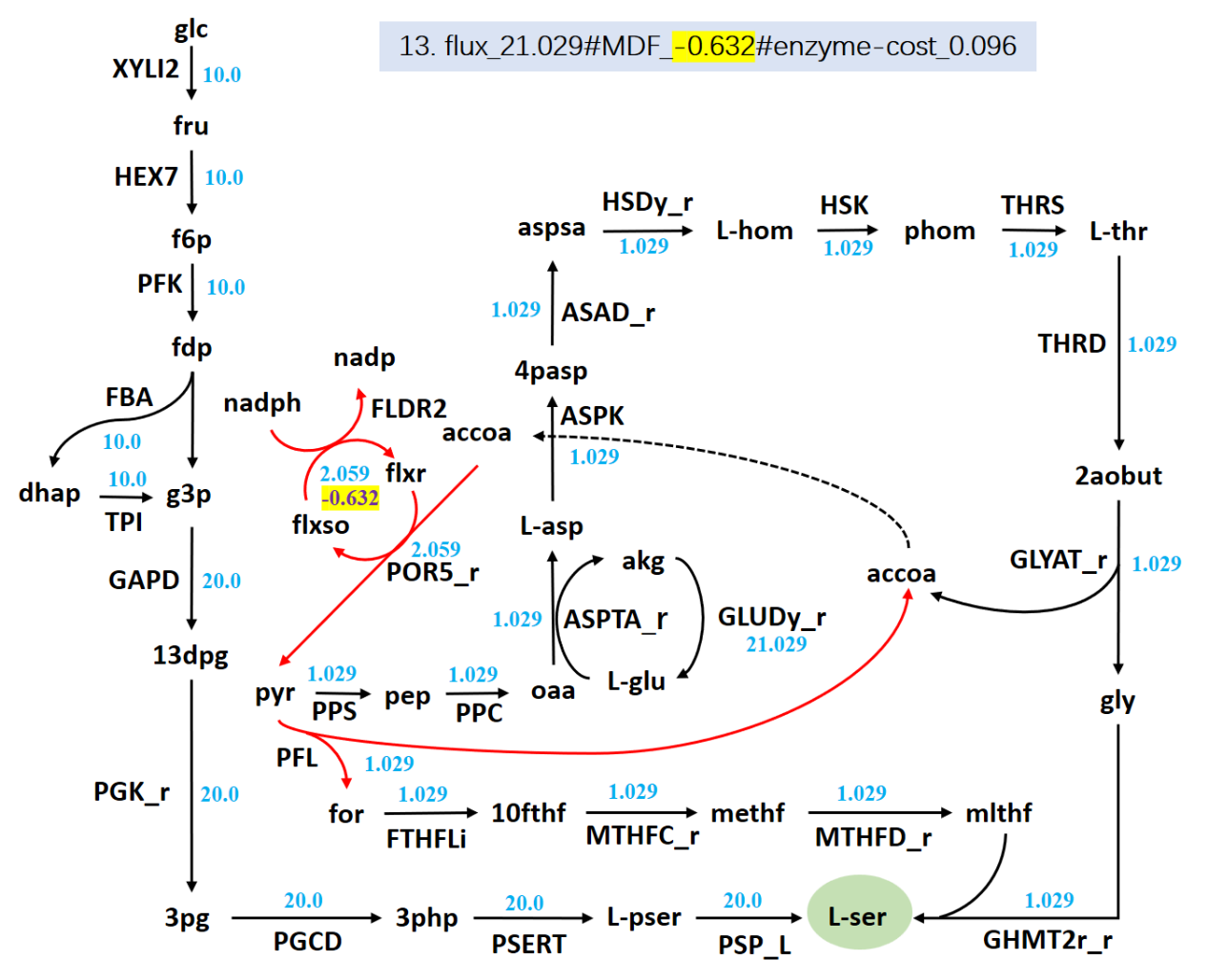

**Figure S1.** The 13 l-serine synthesis pathways point-to-point corresponding to Fig. 1 and table S1. Shown are: the thermodynamic bottleneck reaction (red arrow; since their MDF values are all the same, only one value is listed and marked with yellow background.); The unit of the flux value is mmol/gDW/h (blue); and the unit of the maximum thermodynamic driving force is in kJ/mol; the unit of the enzyme cost is mg/gDW (it’s up bound is 0.13 mg/gDW).

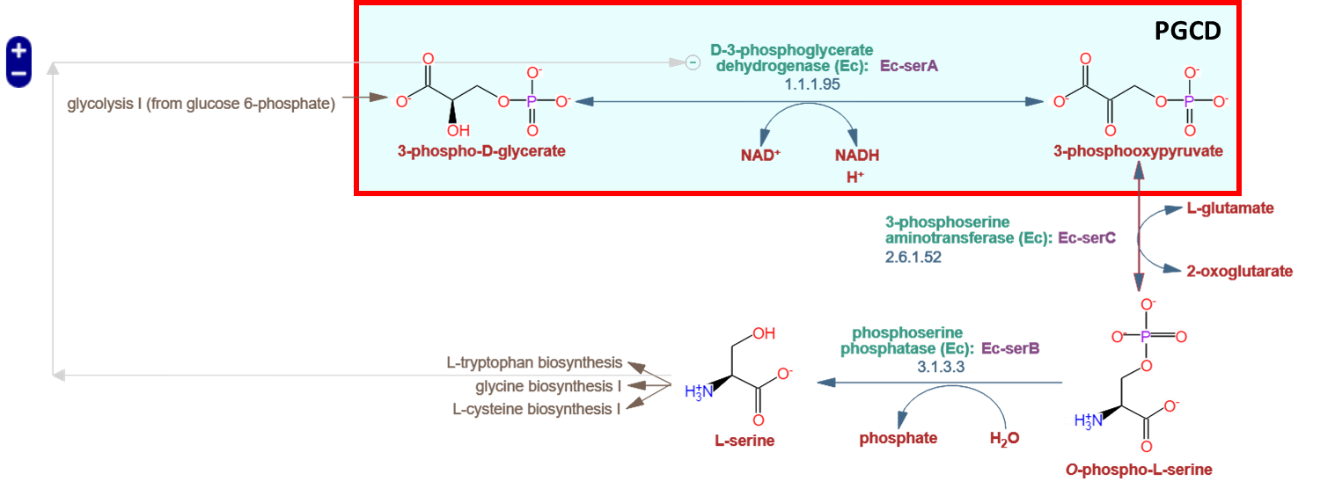

**Figure S2** l-serine synthesis pathway in *E. coli*. The PGCD reaction was marked by red box; the base-map was downloaded from the MetaCyc database (<https://metacyc.org/META/NEW-IMAGE?type=PATHWAY&object=SERSYN-PWY&detail-level=3>).

**Table S2** Three distributed bottleneck reactions causing thermodynamics infeasible

| **No.** | **Reaction** | **Reaction equation** | **max*Df_i_*** |
| --- | --- | --- | --- |
|  | **ID** |  | **kJ/mol** |
| 1 | PFL | coa_c + pyr_c --> accoa_c + for_c | 75.86 |
| 2 | FLDR2 | 2.0 flxso_c + nadph_c --> 2.0 flxr_c + h_c + nadp_c | 45.00 |
| 3 | POR5_r | accoa_c + co2_c + 2.0 flxr_c + h_c --> coa_c + 2.0 flxso_c + pyr_c | 68.55 |

**Table S3** Bottleneck reactions of l-serine synthesis pathways after reaction combination.

| **Turning** | **Max flux** | **MDF** | **Bottleneck** | **Reaction equation** |
| --- | --- | --- | --- | --- |
| **point** | **(mmol/gDW/h)** | **(kJ/mol)** | **reaction** |  |
| 1 | 10.00 | 15.77 | PGK_r | 13dpg_c + adp_c --> 3pg_c + atp_c |
|  |  |  | GAPD | g3p_c + nad_c + pi_c --> 13dpg_c + h_c + nadh_c |
| 2 | 13.81 | 9.86 | ACONTa | cit_c --> acon_C_c + h2o_c |
|  |  |  | ACONTb | acon_C_c + h2o_c --> icit_c |
| 3 | 16.06 | 7.64 | ADK1 | amp_c + atp_c --> 2.0 adp_c |
| 4 | 16.57 | 6.64 | TPI | dhap_c --> g3p_c |
|  |  |  | GAPD | g3p_c + nad_c + pi_c --> 13dpg_c + h_c + nadh_c |
| 5 | 18.77 | 5.54 | TPI | dhap_c --> g3p_c |
|  |  |  | GAPD | g3p_c + nad_c + pi_c --> 13dpg_c + h_c + nadh_c |
|  |  |  | FBA | fdp_c --> dhap_c + g3p_c |

**Table S4** The ***∆_r_G_i_′⁰*** information of newly added overall reaction PFPOr

| No. | Reaction ID | Reaction equation | ***∆_r_G_i_′⁰*** | **Notes** |
| --- | --- | --- | --- | --- |
|  |  |  | **kJ/mol** |  |
| **1** | PFL | coa_c + pyr_c --> accoa_c + for_c | -21.2 ± 3 | Original; Shut |
| **2** | FLDR2 | 2.0 flxso_c + nadph_c --> 2.0 flxr_c + h_c + nadp_c | 15.6 ± 14.9 | Original; Shut |
| **3** | POR5_r | accoa_c + co2_c + 2.0 flxr_c + h_c --> coa_c + 2.0 flxso_c + pyr_c | 27.1 ± 16.7 | Original; Shut |
| **4** | PFPOr | co2_c + nadph_c --> for_c + nadp_c | 21.5 ± 22.6 | Newly; added |

**Table S5** The enzyme related parameters of PFPOr

| **No.** | **Reaction** | **Reaction equation** | ***k*_cat_** | **MW** | |
| --- | --- | --- | --- | --- | --- |
|  | **ID** |  | **h^-1^** | **kDa^-1^** | |
| **1** | PFL | coa_c + pyr_c --> accoa_c + for_c | 204410.7 | | 113.56 |
| **2** | FLDR2 | 2.0 flxso_c + nadph_c --> 2.0 flxr_c + h_c + nadp_c | 47286.67 | | 47.45 |
| **3** | POR5_r | accoa_c + co2_c + 2.0 flxr_c + h_c --> coa_c + 2.0 flxso_c + pyr_c | 52980.57 | | 148.52 |
| 4 | PFPOr | co2_c + nadph_c --> for_c + nadp_c | 47286.67 | | 309.53 |

**Table S6** The MDF level of reaction PFPOr.

| **Reaction ID** | **Reaction equation** | **max*Df_i_*** |
| --- | --- | --- |
|  |  | **kJ/mol** |
| PFPOr | co2_c + nadph_c --> for_c + nadp_c | -1.897 |

**Table S7** Bottleneck reactions of anaerobic l-serine synthesis pathways

| **Turning** | **Max flux** | **MDF** | **Bottleneck** | **Reaction equation** |
| --- | --- | --- | --- | --- |
| **point** | **(mmol/gDW/h)** | **(kJ/mol)** | **reaction** |  |
| 1 | 2.069 | 7.422 | PGK_r | 13dpg_c + adp_c --> 3pg_c + atp_c |
|  |  |  | PGM_r | 3pg_c --> 2pg_c |
|  |  |  | GAPD | g3p_c + nad_c + pi_c --> 13dpg_c + h_c + nadh_c |
|  |  |  | ENO | 2pg_c --> h2o_c + pep_c |
|  |  |  | ASPK | asp__L_c + atp_c --> 4pasp_c + adp_c |
|  |  |  | ASAD_r | 4pasp_c + h_c + nadph_c --> aspsa_c + nadp_c + pi_c |
| 2 | 2.963 | 5.208 | PGK_r | 13dpg_c + adp_c --> 3pg_c + atp_c |
|  |  |  | PGM_r | 3pg_c --> 2pg_c |
|  |  |  | GAPD | g3p_c + nad_c + pi_c --> 13dpg_c + h_c + nadh_c |
|  |  |  | ENO | 2pg_c --> h2o_c + pep_c |
|  |  |  | TPI | dhap_c --> g3p_c |
| 3 | 4.706 | -1.897 | PFPOr | co2_c + nadph_c --> for_c + nadp_c |

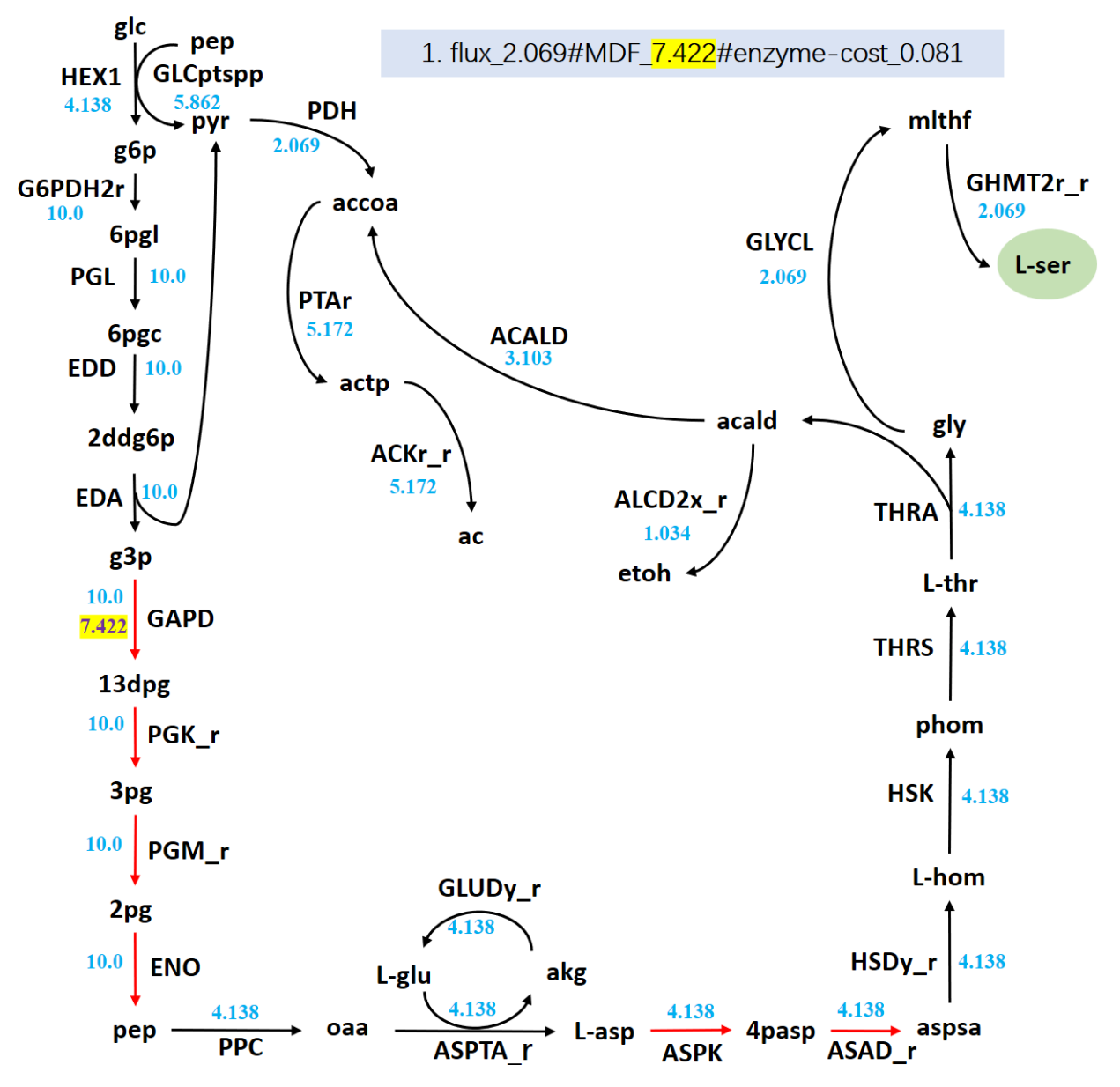

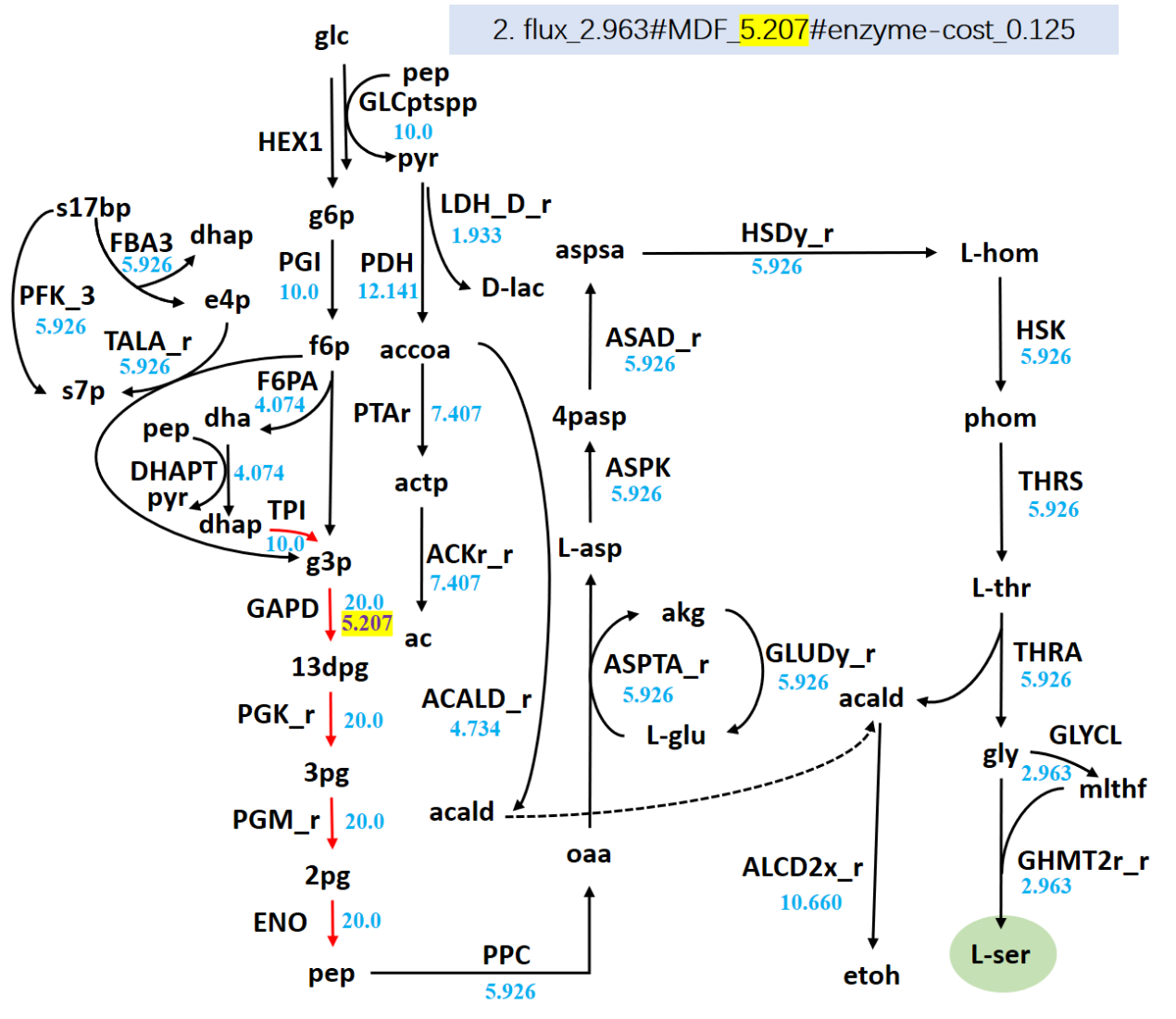

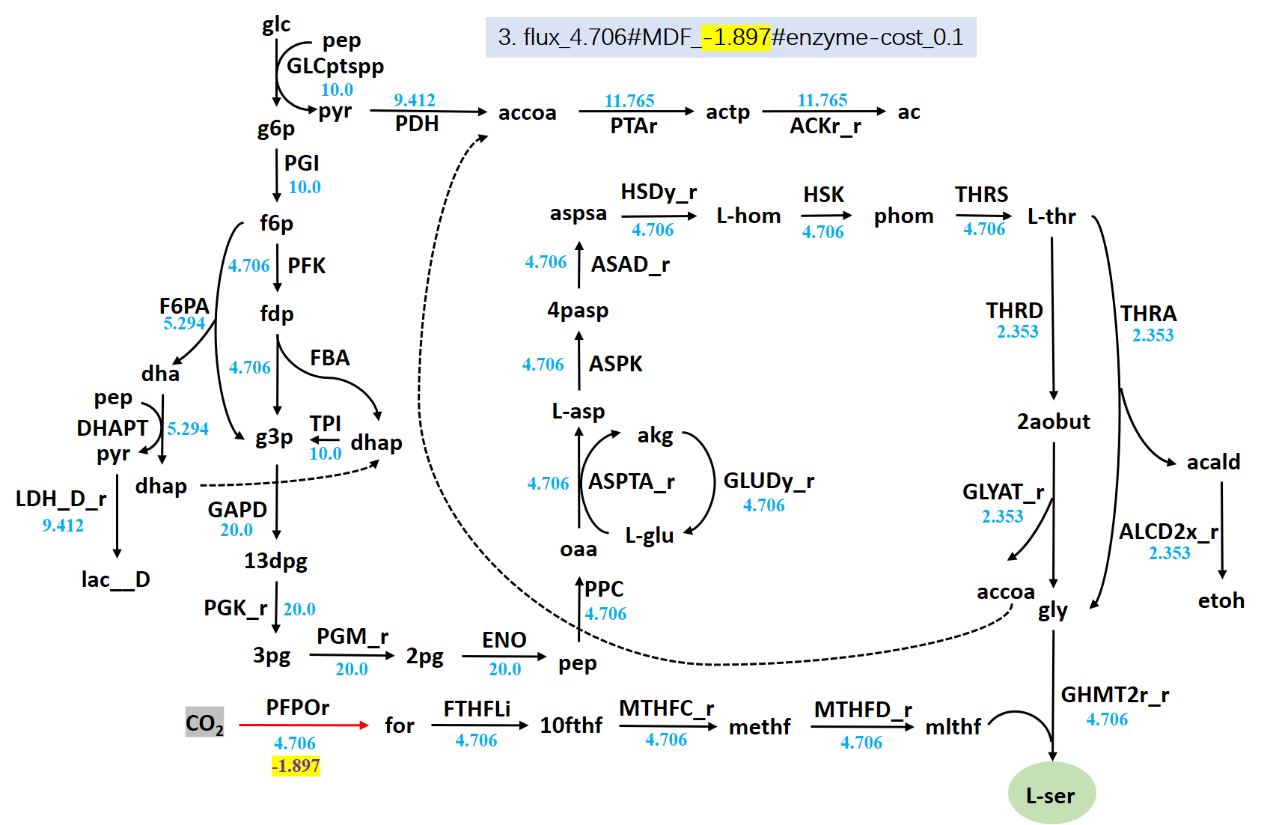

**Figure S3** The 3 l-serine synthesis pathways point-to-point corresponding to table S5. **Table S8** The concentration variability of metabolites in reaction PFL, FLDR2 and POR5_r

| **Metabolite** | **min_ln(conc)** | **max_ln(conc)** | **min_conc.**  **μΜ** | **max_conc.**  **μΜ** | **Notes** |
| --- | --- | --- | --- | --- | --- |
| accoa_c | -6.0432 | -3.9120 | 2.3740 | 20.0000 | / |
| pyr_c | -14.5087 | -12.3775 | 0.0005 | 0.0042 | / |
| nadp_c | -14.5087 | -6.2146 | 0.0005 | 2.0000 | / |
| nadph_c | -12.2061 | -3.9120 | 0.0050 | 20.0000 | / |
| for_c | -14.5087 | -14.5087 | 0.0005 | 0.0005 | lower bound |
| flxr_c | -14.5087 | -5.6625 | 0.0005 | 3.4737 | / |
| co2_c | -9.2103 | -9.2103 | 0.1000 | 0.1000 | up bound |
| flxso_c | -12.7581 | -3.9120 | 0.0029 | 20.0000 | / |
| coa_c | -14.5087 | -12.3775 | 0.0005 | 0.0042 | / |

**Table S9**  The concentration variability of metabolites in overall reaction PFPOr

| **Metabolite** | **min_ln(conc)** | **max_ln(conc)** | **min_conc.**  **μΜ** | **max_conc.**  **μΜ** | **Notes** |
| --- | --- | --- | --- | --- | --- |
| co2_c | -9.2103 | -9.2103 | 0.1000 | 0.1000 | up bound |
| nadph_c | -12.2061 | -3.9120 | 0.0050 | 20.0000 | / |
| nadp_c | -14.5087 | -6.2146 | 0.0005 | 2.0000 | / |
| for_c | -14.5087 | -14.5087 | 0.0005 | 0.0005 | lower bound |

**Table S10** Thermodynamic bottleneck reaction(s) in the L-Trp synthesis pathways predicted by the uncorrected EcoETM

| **Turning** | **Max flux** | **MDF** | **Bottleneck** | **Reaction equation** |
| --- | --- | --- | --- | --- |
| **point** | **(mmol/gDW/h)** | **(kJ/mol)** | **reaction** |  |
| 1 | 1.74 | 9.86 | ACONTa | cit_c --> acon_C_c + h2o_c |
|  |  |  | ACONTb | acon_C_c + h2o_c --> icit_c |
| 2 | 1.78 | 9.82 | TKT2_r | f6p_c + g3p_c --> e4p_c + xu5p__D_c |
|  |  |  | RPI_r | ru5p__D_c --> r5p_c |
|  |  |  | TALA | g3p_c + s7p_c --> e4p_c + f6p_c |
|  |  |  | TKT1 | r5p_c + xu5p__D_c --> g3p_c + s7p_c |
|  |  |  | EDA | 2ddg6p_c --> g3p_c + pyr_c |
|  |  |  | TRPS3 | 3ig3p_c --> g3p_c + indole_c |
| 3 | 1.79 | 9.75 | TPI_r | g3p_c --> dhap_c |
|  |  |  | FBA_r | dhap_c + g3p_c --> fdp_c |
|  |  |  | TRPS3 | 3ig3p_c --> g3p_c + indole_c |
|  |  |  | EDA | 2ddg6p_c --> g3p_c + pyr_c |
|  |  |  | TRPAS2_r | indole_c + nh4_c + pyr_c --> h2o_c + trp__L_c |
| 4 | 1.91 | 9.39 | PRPPS | atp_c + r5p_c --> amp_c + h_c + prpp_c |
|  |  |  | RPI_r | ru5p__D_c --> r5p_c |
|  |  |  | RPE_r | xu5p__D_c --> ru5p__D_c |
|  |  |  | TKT2_r | f6p_c + g3p_c --> e4p_c + xu5p__D_c |
|  |  |  | TRPS3 | 3ig3p_c --> g3p_c + indole_c |
|  |  |  | EDA | 2ddg6p_c --> g3p_c + pyr_c |
|  |  |  | TRPAS2_r | indole_c + nh4_c + pyr_c --> h2o_c + trp__L_c |
| 5 | 2.17 | 8.78 | RPI_r | ru5p__D_c --> r5p_c |
|  |  |  | PPM_r | r5p_c --> r1p_c |
| 6 | 2.87 | 7.64 | ADK1 | amp_c + atp_c <=> 2.0 adp_c |
| 7 | 3.66 | 7.16 | PGK_r | 13dpg_c + adp_c --> 3pg_c + atp_c |
|  |  |  | TRPS3 | 3ig3p_c --> g3p_c + indole_c |
|  |  |  | PGM_r | 3pg_c --> 2pg_c |
|  |  |  | TRPAS2_r | indole_c + nh4_c + pyr_c --> h2o_c + trp__L_c |
|  |  |  | GAPD | g3p_c + nad_c + pi_c --> 13dpg_c + h_c + nadh_c |
|  |  |  | ENO | 2pg_c --> h2o_c + pep_c |
| 8 | 3.82 | 6.28 | PGK_r | 13dpg_c + adp_c --> 3pg_c + atp_c |
|  |  |  | PGM_r | 3pg_c --> 2pg_c |
|  |  |  | GAPD | g3p_c + nad_c + pi_c --> 13dpg_c + h_c + nadh_c |
|  |  |  | ENO | 2pg_c --> h2o_c + pep_c |
|  |  |  | TRPS3 | 3ig3p_c --> g3p_c + indole_c |
|  |  |  | TRPAS2_r | indole_c + nh4_c + pyr_c --> h2o_c + trp__L_c |
| 9 | 3.85 | 5.21 | PGK_r | 13dpg_c + adp_c --> 3pg_c + atp_c |
|  |  |  | PGM_r | 3pg_c --> 2pg_c |
|  |  |  | GAPD | g3p_c + nad_c + pi_c --> 13dpg_c + h_c + nadh_c |
|  |  |  | ENO | 2pg_c --> h2o_c + pep_c |
|  |  |  | TPI | dhap_c --> g3p_c |
| 10 | 4.13 | 4.77 | PGK_r | 13dpg_c + adp_c --> 3pg_c + atp_c |
|  |  |  | PGM_r | 3pg_c --> 2pg_c |
|  |  |  | GAPD | g3p_c + nad_c + pi_c --> 13dpg_c + h_c + nadh_c |
|  |  |  | ENO | 2pg_c --> h2o_c + pep_c |
|  |  |  | TPI | dhap_c --> g3p_c |
|  |  |  | FBA | fdp_c --> dhap_c + g3p_c |

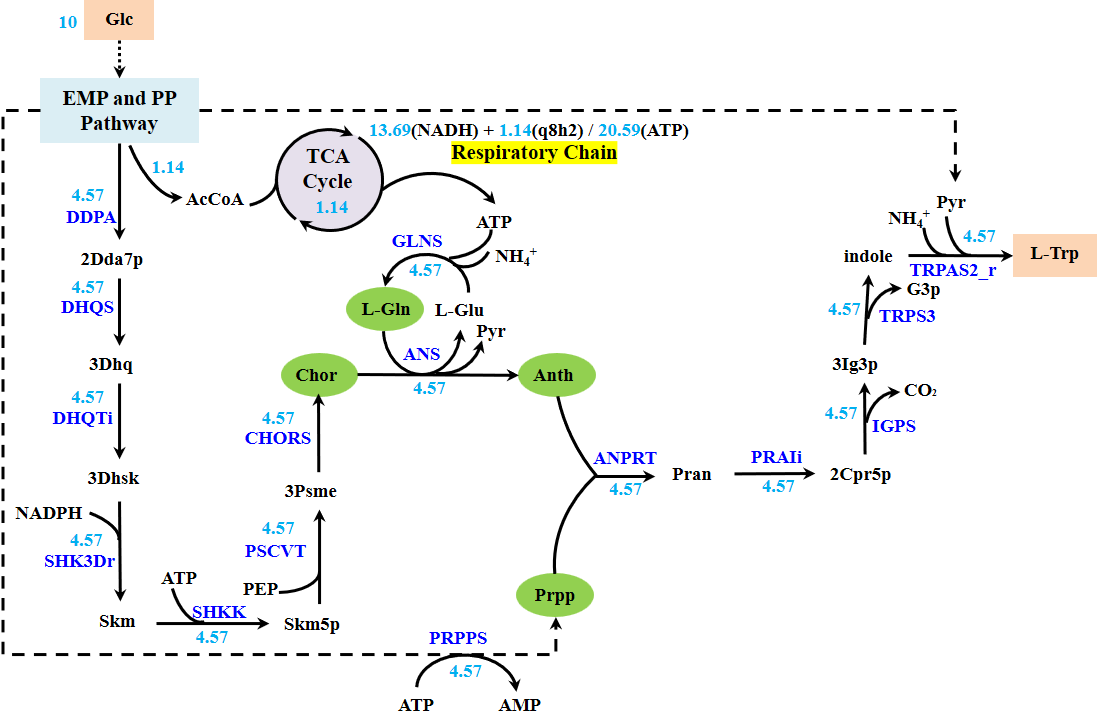

**Figure S4** L-Trp synthesis pathway in *E. coli* predicted by *i*ML1515.

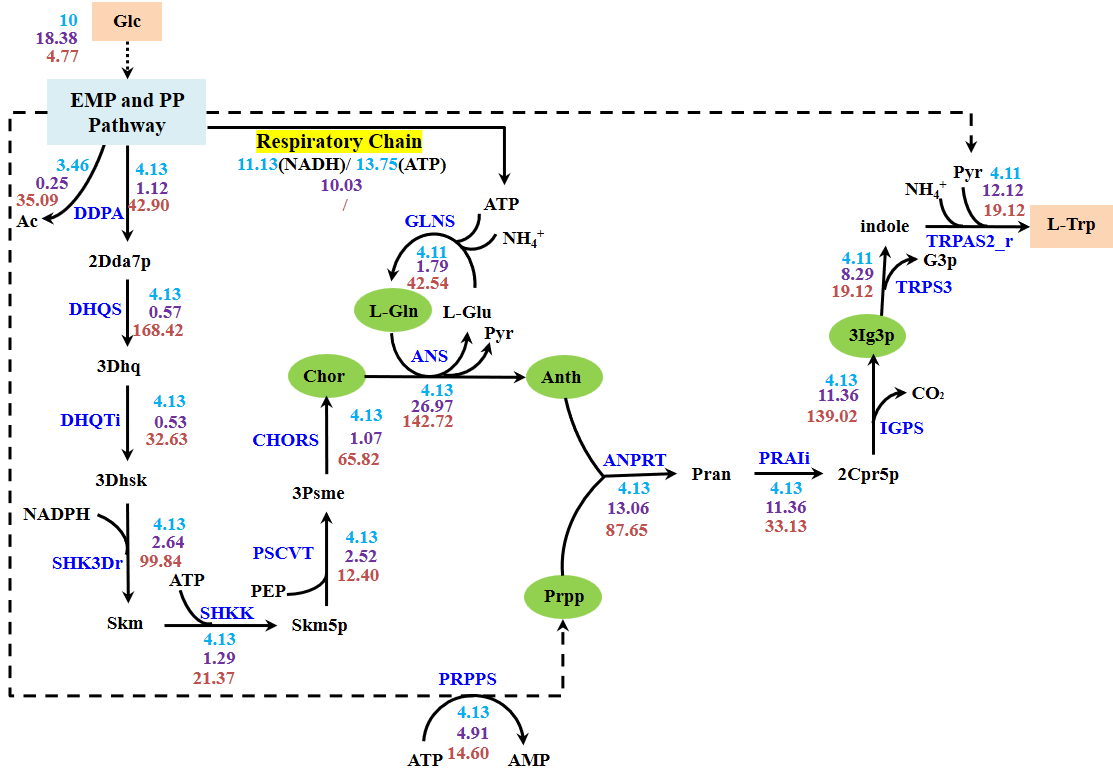

**Figure S5** l-Trp synthesis pathway in *E. coli* predicted by uncorrected EcoETM.

**Table S11** Thermodynamic bottleneck reaction(s) in the L-Trp synthesis processes predicted by the corrected EcoETM

| **Turning** | **Max flux** | **MDF** | **Bottleneck** | **Reaction equation** |
| --- | --- | --- | --- | --- |
| **point** | **(mmol/gDW/h)** | **(kJ/mol)** | **reaction(s)** |  |
| 1 | 1.29 | 17.42 | PRPPS | atp_c + r5p_c --> amp_c + h_c + prpp_c |
|  |  |  | RPI_r | ru5p__D_c --> r5p_c |
| 2 | 1.30 | 17.21 | RPI_r | ru5p__D_c --> r5p_c |
|  |  |  | TPI_r | g3p_c --> dhap_c |
|  |  |  | RPE | ru5p__D_c --> xu5p__D_c |
|  |  |  | PFK_3 | atp_c + s7p_c --> adp_c + h_c + s17bp_c |
|  |  |  | TKT1 | r5p_c + xu5p__D_c --> g3p_c + s7p_c |
| 3 | 1.38 | 16.76 | RPI_r | ru5p__D_c --> r5p_c |
|  |  |  | TKT2_r | f6p_c + g3p_c --> e4p_c + xu5p__D_c |
|  |  |  | TKT1 | r5p_c + xu5p__D_c --> g3p_c + s7p_c |
|  |  |  | PFK_3 | atp_c + s7p_c --> adp_c + h_c + s17bp_c |
| 4 | 1.40 | 16.30 | PGL | 6pgl_c + h2o_c --> 6pgc_c + h_c |
|  |  |  | G6PDH2r | g6p_c + nadp_c --> 6pgl_c + h_c + nadph_c |
|  |  |  | GND | 6pgc_c + nadp_c --> co2_c + nadph_c + ru5p__D_c |
|  |  |  | TKT2_r | f6p_c + g3p_c --> e4p_c + xu5p__D_c |
|  |  |  | RPI_r | ru5p__D_c --> r5p_c |
|  |  |  | TKT1 | r5p_c + xu5p__D_c --> g3p_c + s7p_c |
| 5 | 1.42 | 14.87 | ASPK | asp__L_c + atp_c <=> 4pasp_c + adp_c |
| 6 | 1.90 | 14.71 | HEX7 | atp_c + fru_c --> adp_c + f6p_c + h_c |
|  |  |  | TKT2_r | f6p_c + g3p_c --> e4p_c + xu5p__D_c |
|  |  |  | RPI_r | ru5p__D_c --> r5p_c |
|  |  |  | TKT1 | r5p_c + xu5p__D_c --> g3p_c + s7p_c |
|  |  |  | XYLI2 | glc__D_c --> fru_c |
| 7 | 1.92 | 14.17 | PGK_r | 13dpg_c + adp_c --> 3pg_c + atp_c |
|  |  |  | FBP | fdp_c + h2o_c --> f6p_c + pi_c |
|  |  |  | FBA_r | dhap_c + g3p_c --> fdp_c |
|  |  |  | RPI_r | ru5p__D_c --> r5p_c |
|  |  |  | TKT1 | r5p_c + xu5p__D_c --> g3p_c + s7p_c |
|  |  |  | FBA3 | s17bp_c --> dhap_c + e4p_c |
|  |  |  | PFK_3 | atp_c + s7p_c --> adp_c + h_c + s17bp_c |
|  |  |  | GAPD | 2pg_c --> h2o_c + pep_c |
|  |  |  | TKT2_r | f6p_c + g3p_c --> e4p_c + xu5p__D_c |
| 8 | 1.97 | 12.75 | TALA | g3p_c + s7p_c --> e4p_c + f6p_c |
|  |  |  | TKT2_r | f6p_c + g3p_c --> e4p_c + xu5p__D_c |
|  |  |  | RPI_r | ru5p__D_c --> r5p_c |
|  |  |  | TKT1 | r5p_c + xu5p__D_c --> g3p_c + s7p_c |
| 9 | 2.03 | 11.21 | TALA | g3p_c + s7p_c --> e4p_c + f6p_c |
|  |  |  | PGI | g6p_c --> f6p_c |
|  |  |  | TKT2_r | f6p_c + g3p_c --> e4p_c + xu5p__D_c |
|  |  |  | RPI_r | ru5p__D_c --> r5p_c |
|  |  |  | TKT1 | r5p_c + xu5p__D_c --> g3p_c + s7p_c |
|  |  |  | F6PA | f6p_c --> dha_c + g3p_c |
| 10 | 2.06 | 9.23 | PGI | g6p_c --> f6p_c |
|  |  |  | TKT2_r | f6p_c + g3p_c --> e4p_c + xu5p__D_c |
|  |  |  | RPI_r | ru5p__D_c --> r5p_c |
|  |  |  | RPE_r | xu5p__D_c --> ru5p__D_c |
|  |  |  | PRPPS | atp_c + r5p_c --> amp_c + h_c + prpp_c |
| 11 | 2.07 | 8.92 | PRPPS | atp_c + r5p_c --> amp_c + h_c + prpp_c |
|  |  |  | PGI | g6p_c --> f6p_c |
|  |  |  | TKT2_r | f6p_c + g3p_c --> e4p_c + xu5p__D_c |
|  |  |  | RPI_r | ru5p__D_c --> r5p_c |
|  |  |  | F6PA | f6p_c --> dha_c + g3p_c |
|  |  |  | RPE_r | xu5p__D_c --> ru5p__D_c |
| 12 | 2.34 | 8.78 | RPI_r | ru5p__D_c --> r5p_c |
|  |  |  | PPM_r | r5p_c --> r1p_c |
| 13 | 2.87 | 7.88 | PGK_r | 13dpg_c + adp_c --> 3pg_c + atp_c |
|  |  |  | PGM_r | 3pg_c --> 2pg_c |
|  |  |  | ENO | g3p_c + nad_c + pi_c --> 13dpg_c + h_c + nadh_c |
|  |  |  | GAPD | 2pg_c --> h2o_c + pep_c |
| 14 | 3.10 | 7.64 | ADK1 | amp_c + atp_c <=> 2.0 adp_c |
| 15 | 3.12 | 5.21 | PGK_r | 13dpg_c + adp_c --> 3pg_c + atp_c |
|  |  |  | PGM_r | 3pg_c --> 2pg_c |
|  |  |  | GAPD | g3p_c + nad_c + pi_c --> 13dpg_c + h_c + nadh_c |
|  |  |  | ENO | 2pg_c --> h2o_c + pep_c |
|  |  |  | TPI | dhap_c --> g3p_c |
| 16 | 3.27 | 4.77 | PGK_r | 13dpg_c + adp_c --> 3pg_c + atp_c |
|  |  |  | PGM_r | 3pg_c --> 2pg_c |
|  |  |  | GAPD | g3p_c + nad_c + pi_c --> 13dpg_c + h_c + nadh_c |
|  |  |  | ENO | 2pg_c --> h2o_c + pep_c |
|  |  |  | TPI | dhap_c --> g3p_c |
|  |  |  | FBA | fdp_c --> dhap_c + g3p_c |

**Table S12** Related reactions in the synthesis process of anthranilate

| **No.** | **Reaction ID** | **Reaction equation** | ***∆_r_G_i_′⁰*** | **Notes** |
| --- | --- | --- | --- | --- |
|  |  |  | **kJ/mol** | **Source/Structure** |
| **1** | ADCS | chor_c + gln__L_c --> 4adcho_c + glu__L_c | -16.4±6 | Original; Para- |
| **2** | ADCL | 4adcho_c --> 4abz_c + h_c + pyr_c | -64.1±13.6 | Original; Para- |
| **3** | ANS | chor_c + gln__L_c --> anth_c + glu__L_c + h_c + pyr_c | -80.5±13.1 | Original; Combination |
| **4** | ADCSG | chor_c + gln__L_c --> 2adcho_c + glu__L_c | -16.4±6 | Newly-added; Ortho- |
| **5** | ADCSN | chor_c + nh4_c --> 2adcho_c + h2o_c | -5±5.8 | Newly-added; Ortho- |
| **6** | ADCSG_r | 2adcho_c + glu__L_c --> chor_c + gln__L_c | 16.4±6 | Newly-added; Ortho- |
| **7** | ADCSN_r | 2adcho_c + h2o_c --> chor_c + nh4_c | 5±5.8 | Newly-added; Ortho- |
| 8 | ADCLy | 2adcho_c --> anth_c + h_c + pyr_c | -64.1±13.6 | Newly-added; Ortho- |

**Table S13**  Synthesis reactions and parameters information of anthranilate

| **No.** | **Reaction** | **Reaction equation** | ***k*_cat_** | **MW** | |
| --- | --- | --- | --- | --- | --- |
|  | **ID** |  | **h^-1^** | **kDa^-1^** | |
| **1** | ANS | chor_c + gln__L_c --> anth_c + glu__L_c + h_c + pyr_c | 17526.98 | | 114.36 |
| **2** | ADCSG | chor_c + gln__L_c <=> 2adcho_c + glu__L_c | / | | / |
| **3** | ADCSN | chor_c + nh4_c <=> 2adcho_c + h2o_c | / | | / |
| 4 | ADCLy | 2adcho_c --> anth_c + h_c + pyr_c | 17526.98 | | 114.36 |

**Table S14** Thermodynamic bottleneck reaction(s) in the anthranilate synthesis pathways

| **Turning**  **point** | **Max flux** | | **MDF** | **Bottleneck**  **reaction** | **Reaction equation** |
| --- | --- | --- | --- | --- | --- |
|  | **(mmol/gDW/h)** | | **(kJ/mol)** |  |  |
|  | **L-Gln** | **NH_3_** |  |  |  |
| 1 | 2.861 | / | 19.029 | ACONT | cit_c --> icit_c |
| 2 | 2.869 | / | 17.551 | G6PDH2r | g6p_c + nadp_c --> 6pgl_c + h_c + nadph_c |
|  |  |  |  | GND | 6pgc_c + nadp_c --> co2_c + nadph_c + ru5p__D_c |
|  |  |  |  | PGL | 6pgl_c + h2o_c --> 6pgc_c + h_c |
|  |  |  |  | RPI_reverse | ru5p__D_c --> r5p_c |
|  |  |  |  | TKT1 | r5p_c + xu5p__D_c --> g3p_c + s7p_c |
|  |  |  |  | RPE | ru5p__D_c --> xu5p__D_c |
|  |  |  |  | PFK_3 | atp_c + s7p_c --> adp_c + h_c + s17bp_c |
| 3 | 2.945 | / | 17.205 | TPI_reverse | g3p_c --> dhap_c |
|  |  |  |  | RPI_reverse | ru5p__D_c --> r5p_c |
|  |  |  |  | PFK_3 | atp_c + s7p_c --> adp_c + h_c + s17bp_c |
|  |  |  |  | TKT1 | r5p_c + xu5p__D_c --> g3p_c + s7p_c |
|  |  |  |  | RPE | ru5p__D_c --> xu5p__D_c |
| 4 | 3.412 | / | 16.756 | TKT2_reverse | f6p_c + g3p_c --> e4p_c + xu5p__D_c |
|  |  |  |  | RPI_reverse | ru5p__D_c --> r5p_c |
|  |  |  |  | TKT1 | r5p_c + xu5p__D_c --> g3p_c + s7p_c |
|  |  |  |  | PFK_3 | atp_c + s7p_c --> adp_c + h_c + s17bp_c |
| 5 | 3.461 | / | 16.297 | PGL | 6pgl_c + h2o_c --> 6pgc_c + h_c |
|  |  |  |  | G6PDH2r | g6p_c + nadp_c --> 6pgl_c + h_c + nadph_c |
|  |  |  |  | GND | 6pgc_c + nadp_c --> co2_c + nadph_c + ru5p__D_c |
|  |  |  |  | TKT2_reverse | f6p_c + g3p_c --> e4p_c + xu5p__D_c |
|  |  |  |  | RPI_reverse | ru5p__D_c --> r5p_c |
|  |  |  |  | TKT1 | r5p_c + xu5p__D_c --> g3p_c + s7p_c |
| 6 | 4.036 | / | 14.705 | HEX7 | atp_c + fru_c --> adp_c + f6p_c + h_c |
|  |  |  |  | TKT2_reverse | f6p_c + g3p_c --> e4p_c + xu5p__D_c |
|  |  |  |  | RPI_reverse | ru5p__D_c --> r5p_c |
|  |  |  |  | TKT1 | r5p_c + xu5p__D_c --> g3p_c + s7p_c |
|  |  |  |  | XYLI2 | glc__D_c --> fru_c |
| / | / | 4.138 | 14.514 | ACDSN | chor_c + nh4_c --> 2adcho_c + h2o_c |
| 7 | 4.135 | 4.242 | 14.173 | FBA3 | s17bp_c --> dhap_c + e4p_c |
|  |  |  |  | PGK_reverse | 13dpg_c + adp_c --> 3pg_c + atp_c |
|  |  |  |  | FBP | fdp_c + h2o_c --> f6p_c + pi_c |
|  |  |  |  | TKT2_reverse | f6p_c + g3p_c --> e4p_c + xu5p__D_c |
|  |  |  |  | FBA_reverse | dhap_c + g3p_c --> fdp_c |
|  |  |  |  | RPI_reverse | ru5p__D_c --> r5p_c |
|  |  |  |  | TKT1 | r5p_c + xu5p__D_c --> g3p_c + s7p_c |
|  |  |  |  | PFK_3 | atp_c + s7p_c --> adp_c + h_c + s17bp_c |
|  |  |  |  | GAPD | g3p_c + nad_c + pi_c --> 13dpg_c + h_c + nadh_c |
| 8 | 4.398 | 4.510 | 12.747 | TALA | g3p_c + s7p_c --> e4p_c + f6p_c |
|  |  |  |  | TKT2_reverse | f6p_c + g3p_c --> e4p_c + xu5p__D_c |
|  |  |  |  | RPI_reverse | ru5p__D_c --> r5p_c |
|  |  |  |  | TKT1 | r5p_c + xu5p__D_c --> g3p_c + s7p_c |
| 9 | 4.481 | 4.603 | 11.213 | F6PA | f6p_c --> dha_c + g3p_c |
|  |  |  |  | TALA | g3p_c + s7p_c --> e4p_c + f6p_c |
|  |  |  |  | PGI | g6p_c --> f6p_c |
|  |  |  |  | TKT2_reverse | f6p_c + g3p_c --> e4p_c + xu5p__D_c |
|  |  |  |  | RPI_reverse | ru5p__D_c --> r5p_c |
|  |  |  |  | TKT1 | r5p_c + xu5p__D_c --> g3p_c + s7p_c |
| 10 | 4.516 | 4.643 | 8.437 | TKT2_reverse | f6p_c + g3p_c --> e4p_c + xu5p__D_c |
|  |  |  |  | PGI | g6p_c --> f6p_c |
|  |  |  |  | RPI_reverse | ru5p__D_c --> r5p_c |
|  |  |  |  | TKT1 | r5p_c + xu5p__D_c --> g3p_c + s7p_c |
|  |  |  |  | RPE_reverse | xu5p__D_c --> ru5p__D_c |
| 11 | 4.576 | 4.717 | 8.372 | PGI | g6p_c --> f6p_c |
|  |  |  |  | TKT2_reverse | f6p_c + g3p_c --> e4p_c + xu5p__D_c |
|  |  |  |  | F6PA | f6p_c --> dha_c + g3p_c |
|  |  |  |  | RPI_reverse | ru5p__D_c --> r5p_c |
|  |  |  |  | TKT1 | r5p_c + xu5p__D_c --> g3p_c + s7p_c |
|  |  |  |  | RPE_reverse | xu5p__D_c --> ru5p__D_c |
| 12 | 5.536 | 5.620 | 7.885 | PGK_reverse | 13dpg_c + adp_c --> 3pg_c + atp_c |
|  |  |  |  | PGM_reverse | 3pg_c --> 2pg_c |
|  |  |  |  | GAPD | g3p_c + nad_c + pi_c --> 13dpg_c + h_c + nadh_c |
|  |  |  |  | ENO | 2pg_c --> h2o_c + pep_c |
| 13 | 6.552 | 6.877 | 7.638 | ADK1 | amp_c + atp_c --> 2.0 adp_c |
| 14 | 6.973 | 7.256 | 5.208 | PGK_reverse | 13dpg_c + adp_c --> 3pg_c + atp_c |
|  |  |  |  | TPI | dhap_c --> g3p_c |
|  |  |  |  | PGM_reverse | 3pg_c --> 2pg_c |
|  |  |  |  | GAPD | g3p_c + nad_c + pi_c --> 13dpg_c + h_c + nadh_c |
|  |  |  |  | ENO | 2pg_c --> h2o_c + pep_c |
| 15 | 7.058 | 7.269 | 4.767 | PGM_reverse | 3pg_c --> 2pg_c |
|  |  |  |  | ENO | 2pg_c --> h2o_c + pep_c |
|  |  |  |  | GAPD | g3p_c + nad_c + pi_c --> 13dpg_c + h_c + nadh_c |
|  |  |  |  | PGK_reverse | 13dpg_c + adp_c --> 3pg_c + atp_c |
|  |  |  |  | FBA | fdp_c --> dhap_c + g3p_c |
|  |  |  |  | TPI | dhap_c --> g3p_c |
| 16 | 7.116 | 7.315 | 4.584 | MDH | mal__L_c + nad_c --> h_c + nadh_c + oaa_c |
|  |  |  |  | FUM | fum_c + h2o_c --> mal__L_c |
